# The *Legionella pneumophila* metaeffector Lpg2505 (SusF) regulates SidI-mediated translation inhibition and GDP-dependent glycosyltransferase activity

**DOI:** 10.1101/845313

**Authors:** Ashley M. Joseph, Adrienne E. Pohl, Theodore J. Ball, Troy G. Abram, David K. Johnson, Brian V. Geisbrecht, Stephanie R. Shames

## Abstract

*Legionella pneumophila*, the etiological agent of Legionnaires Disease, employs an arsenal of hundreds of Dot/Icm-translocated effector proteins to facilitate replication within eukaryotic phagocytes. Several effectors, called metaeffectors, function regulate the activity of other Dot/Icm-translocated effectors during infection. The metaeffector Lpg2505 is essential for *L. pneumophila* intracellular replication only when its cognate effector, SidI, is present. SidI is a cytotoxic effector that interacts with the host translation factor eEF1A and potently inhibits eukaryotic protein translation by an unknown mechanism. Here, we evaluated the impact of Lpg2505 on SidI-mediated phenotypes and investigated the mechanism of SidI function. We determined that Lpg2505 binds with nanomolar affinity to SidI and suppresses SidI-mediated inhibition of protein translation. SidI binding to eEF1A and SusF is not mutually exclusive and these proteins bind distinct regions of SidI. We also discovered that SidI possesses GDP-dependent glycosyltransferase activity and that this activity is regulated by Lpg2505. We have therefore renamed Lpg2505, SusF (<u>su</u>ppressor of <u>S</u>idI function). This work reveals novel enzymatic activity for SidI and provides insight into how intracellular replication of *L. pneumophila* is regulated by a metaeffector.

## Introduction

*Legionella pneumophila* is the etiological agent of Legionnaires’ Disease, a severe inflammatory pneumonia that results from uncontrolled bacterial replication within alveolar macrophages. Upon phagocytosis, *L. pneumophila* avoids lysosomal degradation through establishment of an endoplasmic reticulum-derived compartment called the *Legionella*-containing vacuole (LCV) (1). For biogenesis of the LCV and acquisition of nutrients from the host cell, *L. pneumophila* is dependent on a massive arsenal of over 300 individual effector proteins that are translocated directly into the host cell through a Dot/Icm Type IVB secretion system (T4BSS) (2). The cellular functions of the majority of effectors have yet to be elucidated, due in part to their functional redundancy within macrophages (3).

Metaeffectors have emerged as a common theme in *L. pneumophila* pathogenesis and are used by *L. pneumophila* to regulate effector function (4). The first metaeffector described was LubX, which temporally regulates function of its cognate effector SidH by hijacking host ubiquitination machinery to facilitate proteasomal degradation of SidH (5). At least 20 of the over 300 identified *L. pneumophila* effectors are metaeffectors (4) and two of these – SidJ and Lpg2505 – are members of a small group of effectors that are individually important for *L. pneumophila* intracellular replication within macrophages (6–8). SidJ is a glutamylase that covalently modifies and abrogates the function of the SidE family of effector ubiquitin ligases (9–11). Lpg2505 is a metaeffector of unknown function that suppresses toxicity of its cognate effector, SidI (Lpg2504), and is important for intracellular replication only when wild-type *sidI* is expressed (6). This was demonstrated by restoration of *L. pneumophila* Δ*lpg2505* intracellular replication upon either deletion of *sidI* or expression of a non-toxic *sidI* allele (R453P). It was further found that Lpg2505 was sufficient to suppress SidI-mediated toxicity towards the yeast *Saccharomyces cerevisiae* (6). Thus, SidI activity is deleterious to *L. pneumophila* in the absence of Lpg2505, a unique phenotype for a translocated effector.

SidI is one of seven cytotoxic *L. pneumophila* effector proteins that inhibit host cell protein translation (12). Like the majority of *L. pneumophila* effectors, *sidI* alone is dispensable for intracellular replication within macrophages, likely due to functional redundancy with other effectors (12). Other translation inhibiting effectors include Lgt1-3, SidL, LegK4, and Lpg1489 (12–16). Lgt1-3 are glycosyltransferases that inhibit translation by glucosylating eukaryotic elongation factor 1A (eEF1A) at Ser-53 (17–19). LegK4 is an effector kinase that phosphorylates heat-shock protein 70 (Hsp70), thereby reducing its ATPase activity and protein refolding activities (16). SidL and Lpg1489 have been experimentally demonstrated to inhibit host protein synthesis (13, 15), but their modes of action are unknown. Like Lgt1-3, SidI also interacts with eEF1A; however, this interaction is insufficient for translation inhibition (12). SidI also interacts with eEF1Bγ and induces the host heat shock response (12). To date, the mechanism(s) by which SidI functions within the host cell to inhibit host protein synthesis have not been elucidated.

In this study we aimed to discern how Lpg2505 regulates SidI and gain insight into the molecular mechanism of SidI function. We discovered that Lpg2505 and SidI bind with nanomolar affinity and that Lpg2505 suppresses SidI-mediated translation inhibition *in vitro.* Furthermore, SidI interaction with Lpg2505 and eEF1A are not mutually exclusive and these two proteins bind with distinct regions of SidI. Finally, we discovered novel GDP-dependent glycosyltransferase activity for SidI, which is regulated by Lpg2505. For this reason, we have named Lpg2505 as *<u>su</u>ppressor of <u>S</u>idI function* (SusF).

## Results

### SusF and SidI bind with nanomolar affinity

Lpg2505 (SusF) is sufficient to suppress SidI-mediated cytotoxicity and promote *L. pneumophila* intracellular replication (6). To reveal a potential mechanism for SusF-mediated regulation of SidI, we evaluated whether SusF interacts with SidI. Since effectors function within host cells, we investigated whether SidI and SusF interact in the presence of host cell lysates. HEK 293 cells stably producing 3FLAG-eipotpe tagged SusF were generated and we initially attempted to ectopically express GFP-tagged *sidI* within these cells for co-immunoprecipitation. However, wild-type SidI could not be detected, likely due to potent translation inhibition. Thus, we generated recombinant GST-tagged SidI (GST-SidI) and evaluated its ability to interact with 3FLAG-SusF within HEK 293 lysates (see *Materials and Methods*). We found that GST-SidI, but not GST alone, retained 3FLAG-SusF on glutathione-coated beads (**Fig 1A**). Furthermore, GST-SidI - but not GST alone - was retained on Ni-NTA beads coated with recombinant His6-SusF (**Fig 1B**). Thus, SusF interacts with SidI in the presence or absence of mammalian cell lysates.

**Figure 1.**
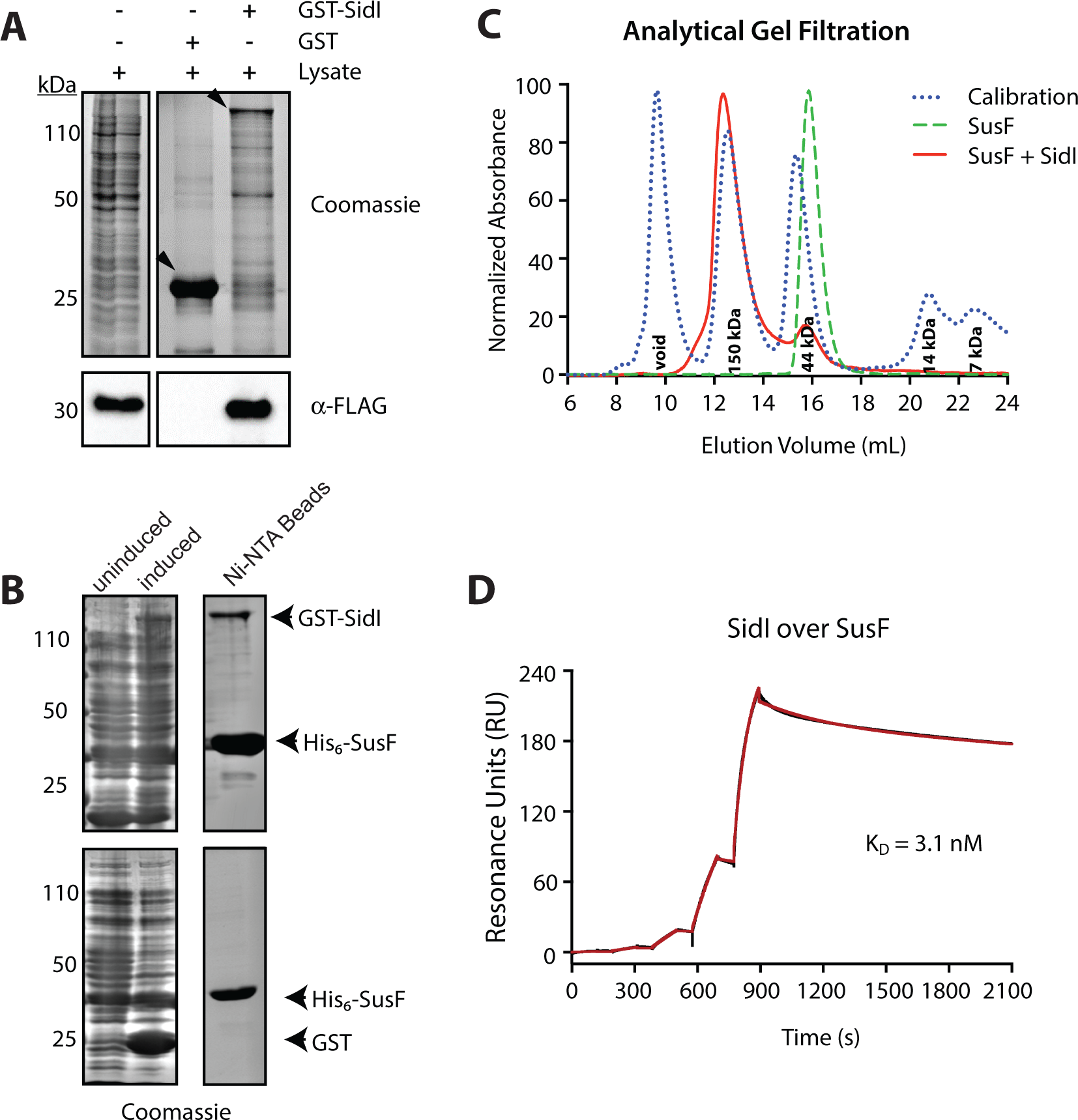
SusF and SidI interact directly with nanomolar affinity. **(A)** Lysates from HEK 293 cells stably expressing 3FLAG-SusF were incubated with glutathione beads coated with either GST or GST-SidI followed by Coomassie staining for total protein and Western blotting for SusF (*α*-FLAG). Arrowheads indicate GST and GST-SidI proteins. **(B)** Lysates from *E. coli* overexpressing either GST or GST-SidI were incubated with Ni-NTA beads coated with His6-SusF followed by SDS-PAGE and Coomassie staining for proteins retained on the beads. Left panel: whole cell lysates from uninduced and induced cultures of *E. coli* expressing GST and GST-SidI proteins; Right panel: proteins retained on Ni-NTA beads (see *Materials and Methods*). **(C)** Chromatograms resulting from analytical scale gel-filtration separation of either SusF alone (green trace) or SusF bound to SidI (red trace). A chromatogram of known size standards is provided for reference (blue trace). **(D)** Binding of His6-SidI to immobilized SusF was assessed by SPR. The reference corrected sensorgram from a single-cycle experiment is shown in black, while the outcome of fitting to a two-state binding model is shown in red. The interaction is described by an apparent KD of 3.1 nM, consisting of two individual steps where kon,1 = 2.7×10^4^ M^-1^s^-1^ and koff,1 = 5.1×10-4 s^-1^ and kon,2 = 2.3×10^-3^ s^-1^ and koff,2 = 4.5×10^-4^ s^-1^, respectively.

To determine if SidI and SusF bind directly, we examined the ability of these proteins to associate with one another throughout the course of sequential column chromatography procedures. SidI and SusF were co-expressed in *Escherichia coli*, purified by Ni-NTA affinity chromatography (see *Materials and Methods*), and the eluted proteins were separated by analytical format size exclusion chromatography. We found that SidI and SusF co-eluted from the column as a species corresponding to a molecular weight of ∼150 kDa, as judged by comparison to a panel of known protein standards (**Fig 1C**). Bands corresponding to both proteins were detected in samples of column fractions that had been separated by SDS-PAGE and analyzed by Coomassie staining (**Fig S1**). Thus, SidI and SusF interact directly and appeared to form a stable complex.

We subsequently used surface plasmon resonance (SPR) to investigate the affinity and kinetics of the SusF-SidI interaction. We immobilized SusF on an SPR surface using random amine chemistry and injected recombinant, purified SidI at increasing concentrations. We employed a single-cycle approach due to difficulty with regenerating the SusF surface following exposure to SidI. We found that the reference-corrected sensorgram could be described fairly well by a Langmuir binding model (KD = 0.89 nM and χ^2^ = 3.26) (**Fig. S2**), but was better fit to a two-state reaction model (KD = 3.1 nM and χ^2^ = 0.79) (**Fig 1D**). A particularly noteworthy feature of the SusF-SidI interaction is its long half-life, with an estimate dissociation rate constant between 2-5×10^-4^ s^-1^. This observation explains, at least in part, the ability of SusF and SidI to remain associated with one another throughout the co-purification procedures described above. Taken together, these data indicate that SusF binds directly to SidI and forms a high-affinity complex with a KD of ∼ 1 nM.

### SusF suppresses SidI-mediated translation inhibition

Based on the high-affinity interaction between SidI and SusF, we hypothesized that SusF represses SidI-mediated translation inhibition. To test this hypothesis, we quantified translation of *Firefly* luciferase mRNA (Luc mRNA) *in vitro* (see *Materials and Methods*). This assay was used previously to demonstrate that ≥ 5 ng of recombinant SidI was sufficient to completely abolish translation (12). We confirmed that purified His6-SidI significantly attenuates protein translation and additionally revealed that purified SusF alone has no impact on translation (**Fig 2A**). We further determined that SusF significantly rescues translation in the presence of SidI (*P*<0.01, **Fig 2B**). Thus, SusF is sufficient to suppress SidI-mediated translation inhibition *in vitro*.

**Figure 2.**
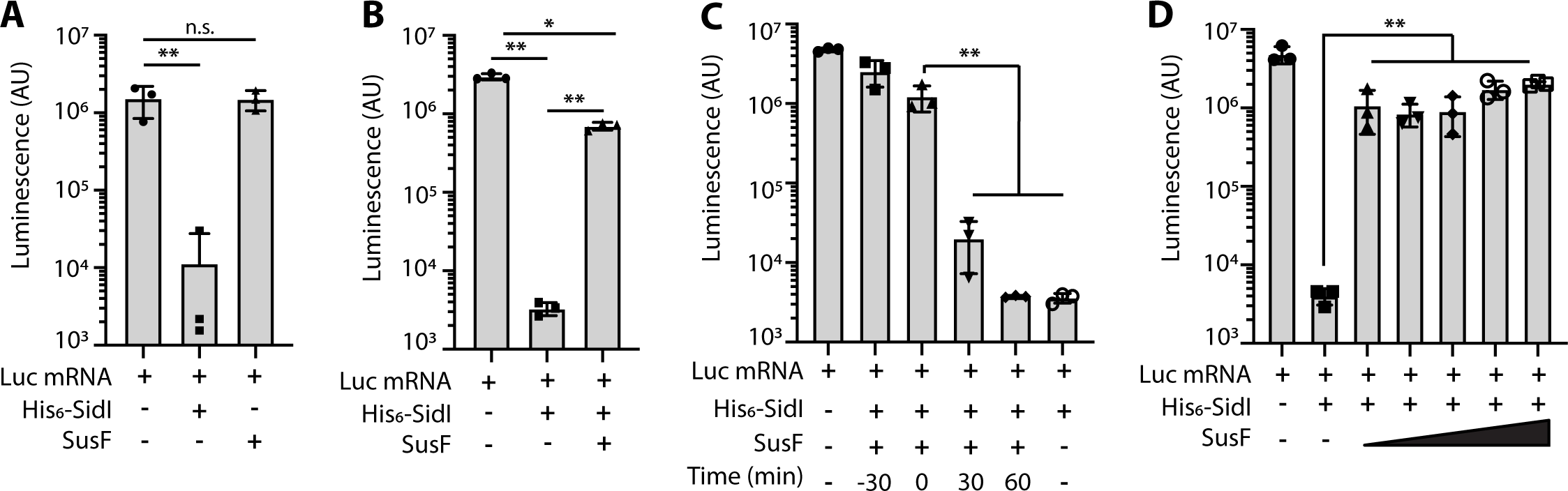
SusF modulates SidI-mediated translation inhibition *in vitro*. Translation of luciferase mRNA was quantified using a rabbit reticulocyte lysate kit. Translation of luciferase in the presence of **(A)** 4 ng His6-SidI or 20 ng SusF; **(B)** His6-SidI alone or 4 ng (37 fmol) His6-SidI and an equimolar amount of SusF (37 fmol); **(C)** His6-SidI alone or His6-SidI and SusF were added to *in vitro* translation reactions at the indicated times (−30 indicates pre-incubation of His6-SidI and SusF for 30 min prior to translation reaction); **(D)** Quantification of *In vitro* translation of Luc mRNA in the presence of His6-SidI alone or His6-SidI and SusF at increasing molar ratios (1:1, 2:1 and 5:1 molar ratios of SusF:His6-SidI). Data are representative of at least two independent experiments and shown as mean ± s.d. of samples in triplicates. Asterisks denote statistical significance by *t*-test (\**P*<0.05, \*\**P*<0.01; n.s., not significant).

Our SPR data demonstrates that the SidI-SusF interaction occurs rapidly. We therefore hypothesized that pre-formation of the SidI-SusF complex would not result in further attenuation of SidI-mediated translation inhibition. To determine whether or not pre-formation of the SidI-SusF complex would enhance SusF-mediated suppression of translation inhibition, we incubated SidI with SusF for 30 min prior to addition to the *in vitro* translation reaction. Pre-formation of the SidI-SusF complex did not further attenuate SidI-mediated translation inhibition (**Fig 2C**; see *Materials and Methods*). Furthermore, SusF was insufficient to reverse SidI-mediated translation inhibition since addition of SusF to the reaction after either 30 min or 60 min did not restore translation (**Fig 2C**). Finally, we evaluated whether suppression of SidI-mediated translation inhibition by SusF was dose dependent. SusF was added in concentrations ranging from equimolar up to a 15-fold molar excess relative to SidI. An equimolar amount of SusF was sufficient to suppress SidI-mediated translation inhibition and translation was not further enhanced by addition of up to a 15-fold molar excess of SusF (**Fig 2D**). Thus, equimolar amounts of SusF are sufficient to suppresses SidI-mediated translation inhibition.

### SusF does not affect the interaction between SidI and eEF1A

A previous report demonstrated that SidI interacts directly with the transcription factor eukaryotic elongation factor 1A (eEF1A) (12). We therefore investigated whether the interactions between SidI and eEF1A or SusF are mutually exclusive. GST-SusF or GST alone were immobilized on glutathione beads and incubated with His6-SidI that was pre-incubated with HEK 293T lysates. We found that eEF1A was retained on the beads only in the presence of SidI and that eEF1A did not impair interaction between SusF and SidI (**Fig 3A**). Subsequently, we asked whether increasing the concentration of SusF would influence the SidI-eEF1A interaction. We incubated GST-SidI with eEF1A (from HEK 293T lysates, see *Materials and Methods*) followed by increasing concentrations of purified recombinant SusF. Despite SusF concentration increasing 100-fold, GST-SidI still retained eEF1A on the beads (**Fig 3B**), suggesting that SidI interacts with SusF and eEF1A simultaneously. The same result was observed when SusF was incubated with GST-SidI prior to HEK 293T lysates (**Fig S3**). Thus, SidI binding to eEF1A and SusF is not mutually exclusive.

**Figure 3.**
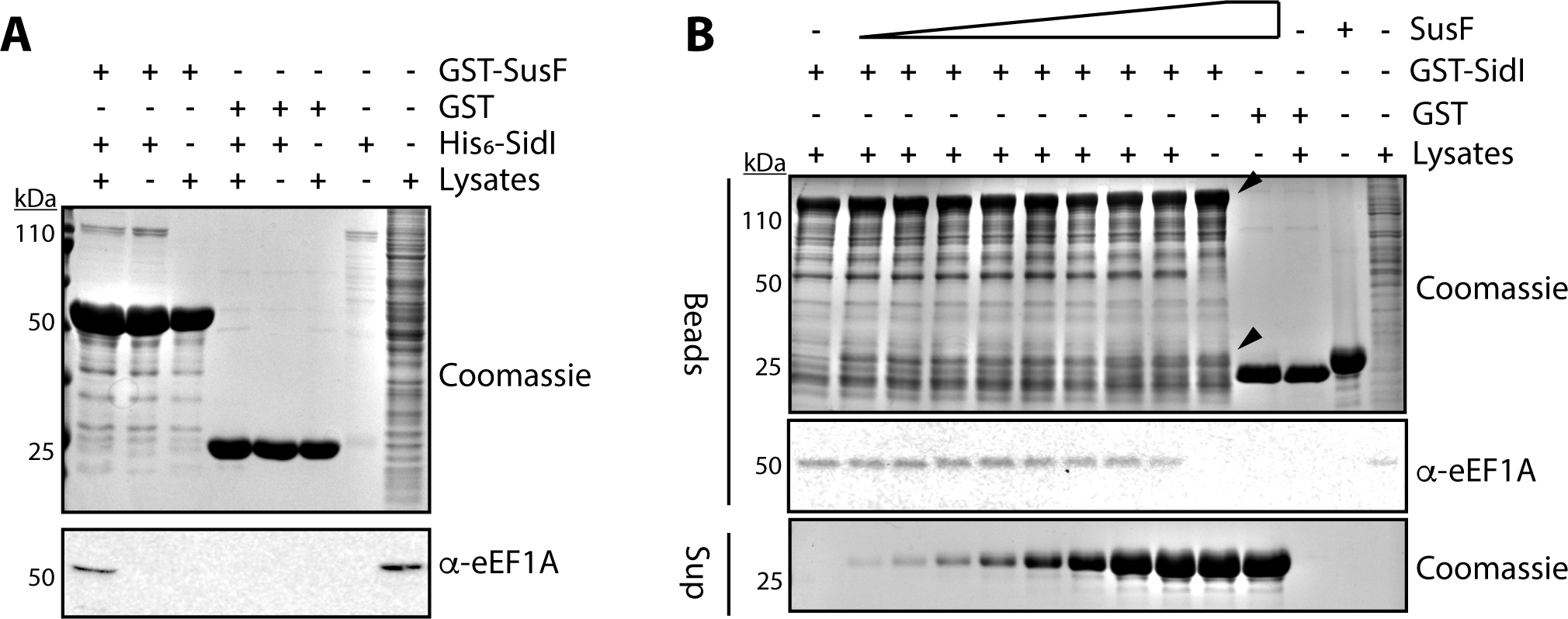
SusF does not impair interaction between SidI and eEF1A. **(A)** GST-SusF or GST alone were immobilized on magnetic glutathione agarose beads followed by incubation with 100 µg purified His6-SidI alone or that had been preincubated with lysates from HEK 293T cells as indicated (Lysates). Proteins were separated by SDS-PAGE and visualized by Coomassie staining or Western blot as indicated. **(B)** Lysates from *E. coli* expressing GST-SidI or GST alone were incubated with magnetic glutathione agarose beads and lysates from HEK 293T followed by 10, to 1000 µg of purified recombinant SusF (shown as increasing amounts in supernatants from beads). Proteins remaining on the beads were separated by SDS-PAGE and visualized by Coomassie stain or Western blot as indicated. GST-SidI (∼130 kDa) and SusF (∼27 kDa) are indicated with arrowheads. Data are representative of at least two independent experiments.

### SusF and eEF1A interact with distinct regions of SidI

Since both SusF and eEF1A are capable of binding SidI simultaneously, we hypothesized that these proteins interact with SidI at distinct sites. SidI has a molecular weight of ∼ 110 kDa and consists of 942 amino acids. Since the structure of SidI has not been solved, we used the RaptorX webserver (20) to predict the domain structure of SidI. Based on the predicted domains (Fig S4), we generated SidI truncations consisting of amino acid residues 1-268 (SidIN), 269-942 (SidIC) and 269-874 (SidICΔ68) (**Fig 4A**). To determine which of these putative SidI domains is involved in interaction with SusF and eEF1A, we immobilized GST-tagged full-length SidI, SidIN, SidIC, SidICΔ68, or GST alone on glutathione beads followed by incubation with lysates from HEK 293 cells stably producing 3FLAG-SusF. Western blot analysis was used to detect 3FLAG-SusF and eEF1A bound to SidI truncation proteins (see *Materials and Methods*). We confirmed that GST-SidI, but not GST alone, was capable of retaining both SusF and eEF1A on the beads. 3FLAG-SusF was further retained on beads coated with GST-SidIN and GST-SidIC but not GST-SidICΔ68, whereas eEF1A was retained by GST-SidIC and GST-SidICΔ68 (**Fig 4B**). To control for the potential influence of 3FLAG-SusF on eEF1A binding to SidI, we repeated this experiment using HEK 293 cells that do not express *3FLAG-susF*. We observed the same pattern of interactions between eEF1A and the C-terminal region of SidI in the absence of 3FLAG-SusF (**Fig S5**). Based on these data, we conclude that SusF interacts with SidI at regions within amino acid residues 1-268 and 874-942 whereas eEF1A interaction with SidI is dependent on amino acid residues 269-874. Together, these data suggest SusF interacts with two regions of SidI that are distinct from the site of SidI interaction with eEF1A.

**Figure 4.**
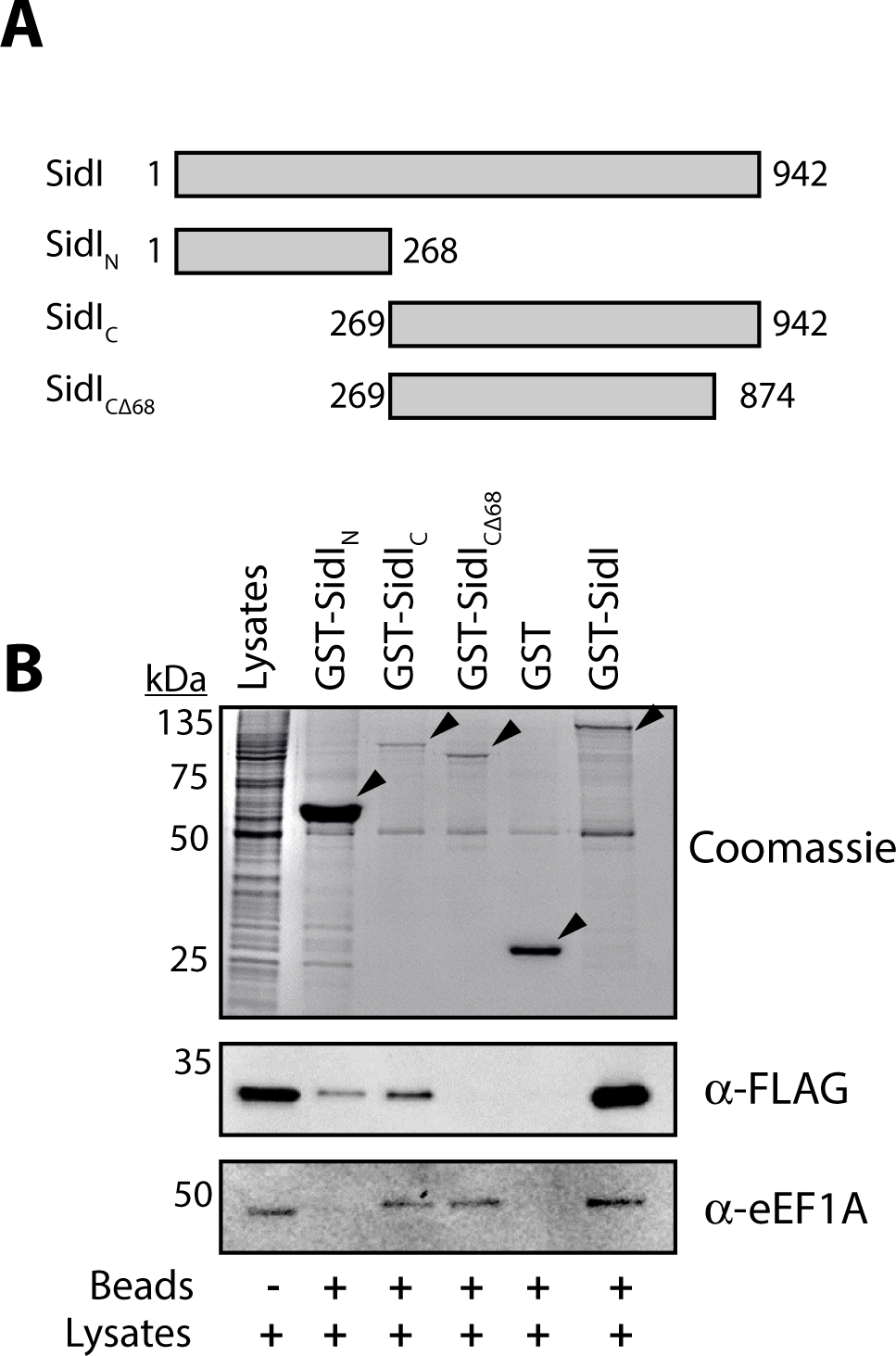
SusF and eEF1A interact with distinct regions of SidI. **(A)** Schematic representation of SidI truncation proteins. **(B)** Lysates from *E. coli* expressing GST-SidI constructs were incubated with glutathione agarose beads followed by washing and incubation with lysates from HEK 293 cells stably expressing 3FLAG-SusF as indicated. Proteins were separated by SDS-PAGE and visualized by Coomassie stain (GST-SidI) or Western blot. Arrowheads indicate fusion proteins. Data are representative of three independent experiments.

### SidI is not dependent on SusF for translocation into host cells

The majority of *L. pneumophila* effector proteins rely on a C-terminal translocation signal for Dot/Icm-mediated translocation into host cells (21, 22). Based on the observed interaction between SusF and the C-terminal 68 amino acid residues of SidI, we hypothesized that SusF may impact Dot/Icm-mediated translocation of SidI. To test this hypothesis, we quantified the export of SidI fused to *Bordetella pertussis* adenylate cyclase (CyaA) (21). When CyaA fusion proteins reach the cytosol of eukaryotic cells they catalyze formation of cAMP, which can be quantified using cAMP-specific ELISA. Based on potent toxicity associated with wild-type SidI, a non-toxic *sidI* allele (R453P; SidIRP) was used for these experiments (6, 12). Expression of CyaA-SusF and CyaA-SidIRP was confirmed using Western blot analysis (**Fig 5A**). Subsequently, we found that cAMP production by host cells was significantly increased following infection with wild-type and Δ*susF*, but not Δ*dotA*, strains of *L. pneumophila* producing CyaA-SidIRP (**Fig 5B**). Translocation of CyaA-RalF was used as a positive control for Dot/Icm-dependent translocation (21) (**Fig 5A, B**). Together, these data demonstrate that SidI is not dependent on SusF for translocation into host cells.

**Figure 5.**
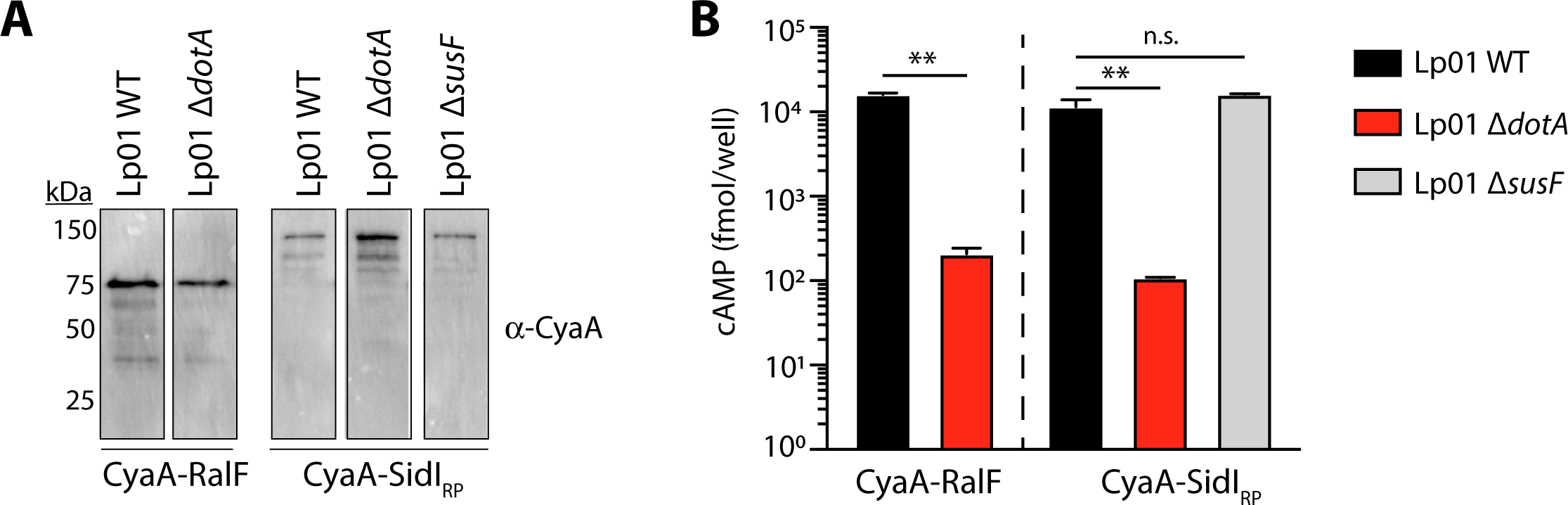
SusF is not required for Dot/Icm translocation of SidI into host cells. **(A)** Western blot showing production of CyaA-RalF (∼76 kDa) and CyaA-SidIRP (∼150 kDa) by the indicated *L. pneumophila* strains. **(B)** Quantification of cAMP extracted from CHO Fc*γ*RII cells infected with the indicated *L. pneumophila* strains. Asterisks denote statistical significance by *t*-test (\*\**P*<0.01). Data are representative of two independent experiments (n.s., not significant).

### Binding to eEF1A is insufficient for SidI-mediated translation inhibition

Subsequently, we investigated the ability of truncated SidI proteins to influence protein translation. At concentrations equivalent to SidI, neither SidIN nor SidICΔ68 were sufficient to inhibit protein translation *in vitro* (**Fig 6A**). Based on the interaction between SidICΔ68 and eEF1A, we also quantified translation in the presence of increasing concentrations of His6-SidICΔ68. We found that His6-SidICΔ68 significantly decreased translation at concentrations up to 25-fold greater than SidI (**Fig 6B**), suggesting that this truncation retains some activity; however, SidICΔ68-mediated translation inhibition is modest in comparison to an equal amount of SidI despite the ability to interact with eEF1A (**Fig 4B**, Fig S5). These data further confirm that interaction with eEF1A is insufficient for SidI-mediated translation inhibition (12).

**Figure 6.**
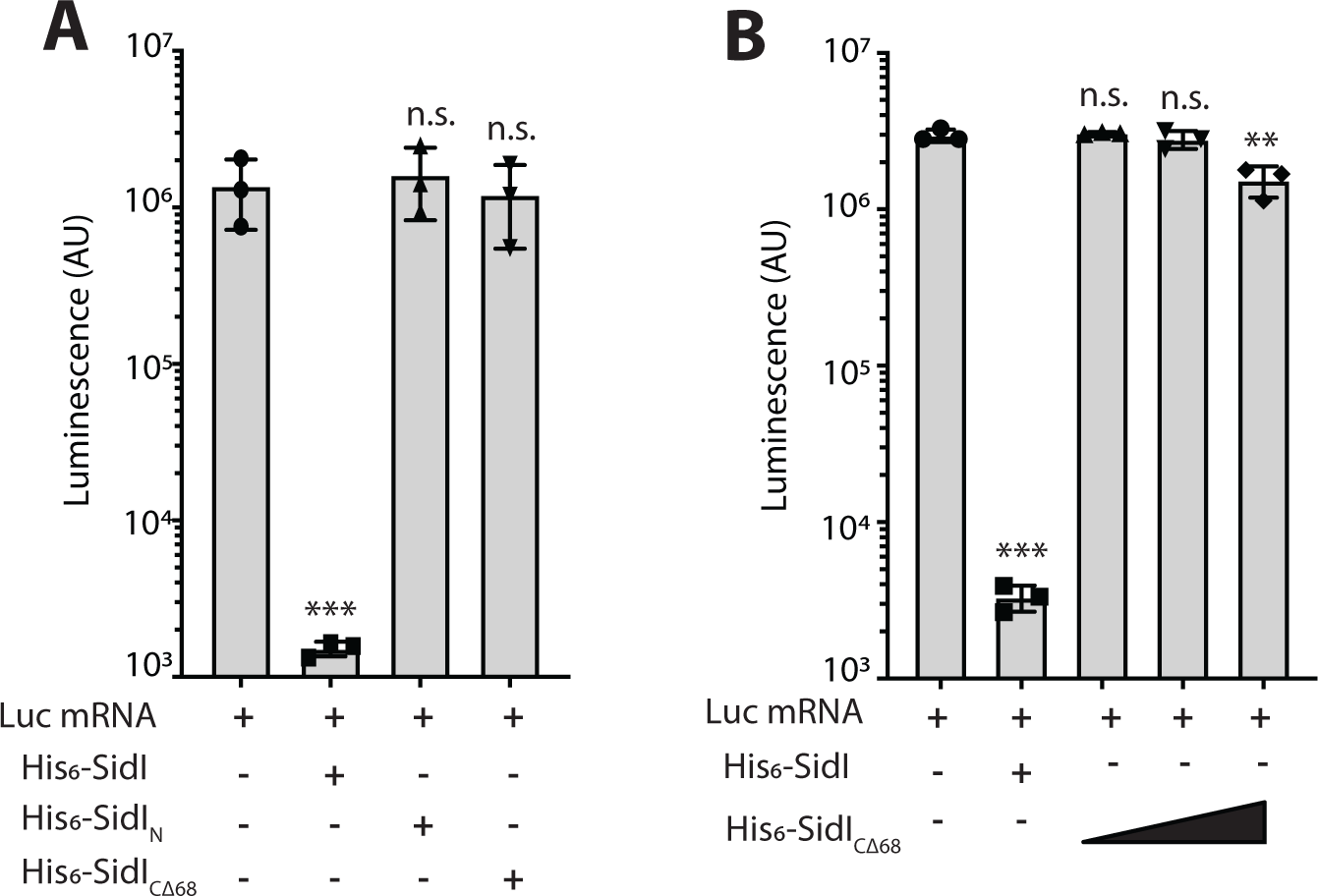
Full-length SidI is required for translation inhibition. Translation of luciferase mRNA quantified using a rabbit reticulocyte lysate kit. **(A)** Translation of luciferase mRNA in the presence of **(A)** 4 ng His6-SidI, His6-SidIN, His6-SidIC, or His6-SidICΔ68 or (**B)** 4 ng of His6-SidI alone or increasing amounts of His6-SidICΔ68 (4 ng, 50 ng, 100 ng). Asterisks denote statistical significance by *t*-test (\*\**P*<0.01). Data are representative of at least two independent experiments.

### SidI is a GDP-dependent glycosyltransferase

Like many other bacterial effectors, the primary amino acid sequence of SidI does not have obviously conserved motifs. Therefore, to gain insight into the putative function of SidI, we used a variety of computational methods to predict the structure and function of SidI. Though very low primary sequence identity was present in the templates used, the HHPred webserver (23), the Phyre2 webserver (24), the Raptor-X webserver (20), and the I-TASSER webserver (25–27) all produced models of various lengths, but with the same fold for overlapping regions. A DALI search (28) for structural homologs against each of these models revealed that amino acid residues 368-868 of SidI have predicted structural homology to multiple bacterial and eukaryotic glycosyltransferases with GT-B folds, including PimB, a GDP-mannose-dependent mannosyltransferase (**Table S1**, Fig S6). Orthogonally, the Ginzu domain parser on the Robetta server also predicted the presence of a glycosyltransferase domain (29). Based on the consensus between orthogonal computational approaches, we hypothesized that SidI possesses GDP-dependent glycosyltransferase activity and that GDP-mannose could be a substrate. To test this hypothesis, we utilized a functional luminescence-based assay to quantify cleavage of GDP-mannose (see *Materials and Methods*). As a control, we also evaluated the ability of SusF, which does not have predicted glycosyltransferase activity, to cleave GDP-mannose. We observed high levels of free GDP only following incubation of recombinant His6-SidI with GDP-mannose, suggesting that SidI indeed possesses GDP-dependent glycosyltransferase activity (**Fig 7A**). We further investigated whether SidI glycosyltransferase activity was specific to GDP-sugars and evaluated its ability to cleave UDP-glucose, which is cleaved by Lgt1-3 (18, 30). SidI was capable of cleaving UDP-glucose, but far less efficiently than GDP-mannose (**Fig 7B**), suggesting that GDP-mannose is the preferred substrate. Together, these data suggest that SidI functions specifically as a GDP-dependent glycosyltransferase.

**Figure 7.**
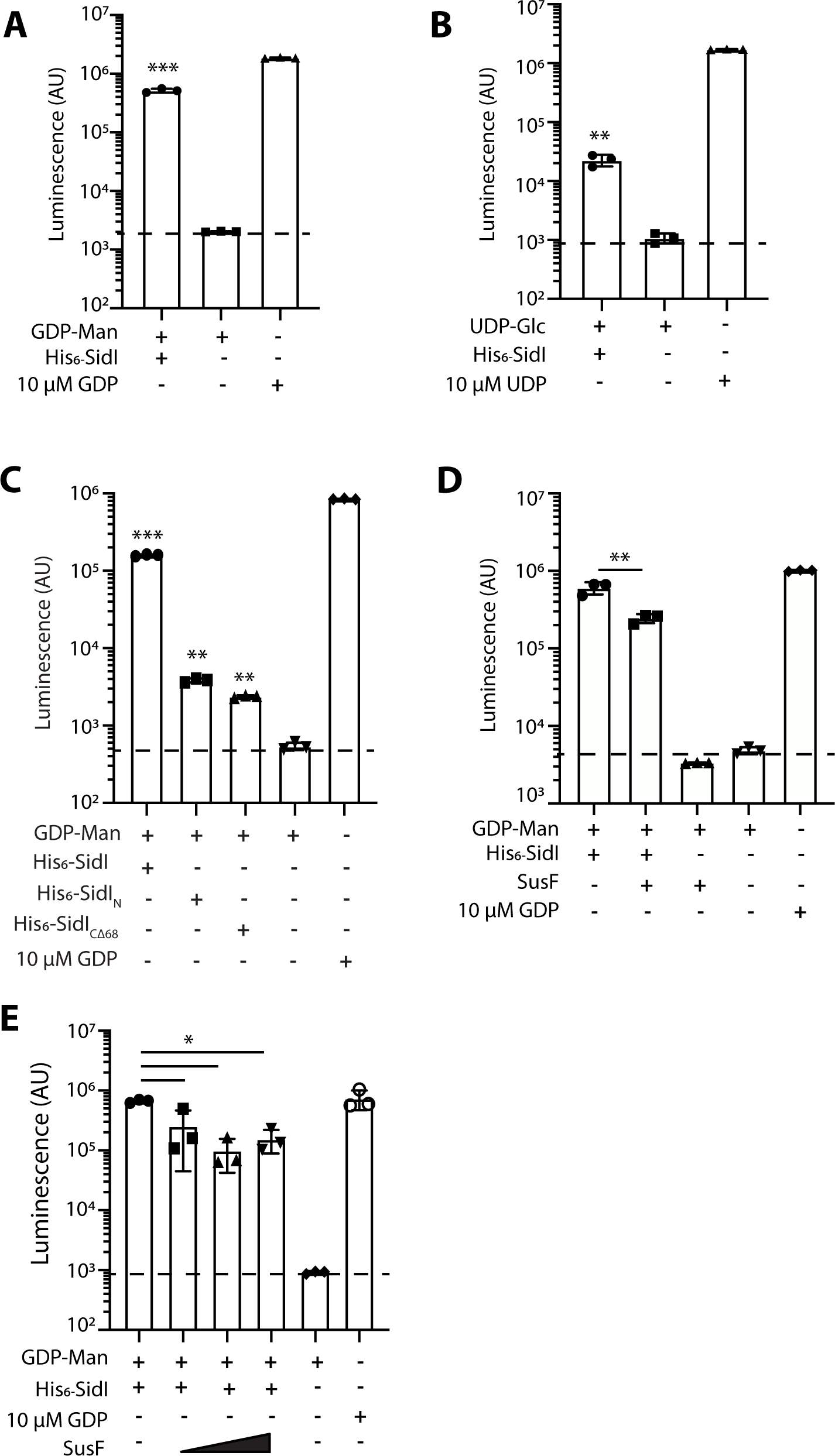
SidI possesses glycosyltransferase activity. **(A)** Quantification of GDP liberated from GDP-mannose in the presence or absence of 5 µg purified His6-SidI. **(B)** Quantification of UDP liberated from UDP-glucose in the presence or absence of 5 µg purified His6-SidI. Quantification of free GDP from GDP-mannose in the presence or absence of **(C)** 5 µg His6-SidI full-length or truncated proteins, **(D)** 5 µg His6-SidI and/or molar equivalent of SusF, or **(E)** increasing molar amount of SusF (1:1, 2:1, or 5:1 molar ratio of SusF:His6-SidI). Asterisks denote statistical significance by *t*-test (\*\**P*<0.01 and \**P*<0.05). Data are representative of at least two independent experiments.

The entire predicted GDP-dependent glycosyltransferase domain of SidI is encoded by amino acids 350-874, which is encompassed within SidICΔ68 truncation (**Fig 4A**). To determine whether SidICΔ68 alone is sufficient to cleave GDP-mannose, we incubated His6-SidICΔ68 with GDP-mannose and quantified generation of free GDP. Full-length SidI and SidIN, which is not predicted to possess glycosyltransferase activity, were included as controls. Neither SidIN nor SidICΔ68 were sufficient to cleave GDP-mannose (**Fig 7C**), suggesting that SidI enzymatic activity is dependent on both the N- and C-terminal domains.

### SidI enzymatic activity is dampened by SusF

Based on the ability of SusF to suppress SidI-mediated translation inhibition, we hypothesized that SusF could regulate SidI-mediated GDP-dependent glycosyltransferase activity. Thus, SidI-mediated GDP-mannose cleavage in the presence of equimolar amounts of SusF was quantified as above. We found that SusF was sufficient to significantly decrease – but not inhibit – SidI GDP-dependent glycosyltransferase activity (\*\**P* < 0.01, **Fig 7D**). We further evaluated whether addition of molar excess of SusF would further decrease SidI glycosyltransferase activity; however, addition of up to a 5-fold molar excess of SusF to SidI was not sufficient to significantly inhibit SidI activity compared to equimolar quantities (**Fig 7E**). Thus, SusF dampens SidI glycosyltransferase activity *in vitro*.

## Discussion

*Legionella pneumophila* is an opportunistic, intracellular pathogen that exploits host cell machinery to promote intracellular replication through the translocation of effector proteins. SidI is one of seven cytotoxic effectors that inhibit eukaryotic protein translation (SidI, SidL, Lgt1-3, LegK4, Lpg1489) (12, 13, 15, 18). Within this family of effectors, SidI is distinct as its regulation by the metaeffector Lpg2505 (SusF) is essential for intracellular replication (6). Our study aimed to explore the role of SusF in regulation of SidI function and elucidate potential mechanisms behind SidI-mediated toxicity. Here we demonstrate that SusF binds directly to SidI with high affinity and modulates both SidI-mediated translation inhibition and novel GDP-dependent glycosyltransferase activity, which has not been previously observed for a bacterial effector. We further demonstrate that SidI is able to simultaneously bind SusF and eEF1A, which interact with distinct regions of SidI. This work is the first to define the enzymatic activity of SidI and the contribution of SusF to SidI-mediated phenotypes.

*L. pneumophila* is reliant on the host cell-derived amino acids for intracellular replication (31). It can therefore be speculated that *L. pneumophila* utilizes multiple effector proteins to halt host protein synthesis at the elongation step in order to facilitate proteasomal degradation of partially folded polypeptides. The translation inhibiting effectors characterized to date have diverse functions. The effector Lgt1 was found to target the host elongation factor eEF1A and homology searches led to the discovery of the orthologous effectors Lgt2-3(19). The Lgt effectors function by glycosylation of eEF1A at Ser-53, which results in blockade of host protein synthesis (32). The Lgts also glycosylate a eukaryotic release factor related protein (eRF3) and the Hsp70 subfamily B suppressor 1 (Hbs1) (33, 34). Based on the function of Lgt1-3, attempts were made to identify SidI-mediated post-translational modification of eEF1A; however, no modifications were discovered (12). Although SidI was previously assumed to lack glycosyltransferase activity (12, 35), we discovered that SidI, like the Lgts, is indeed able to cleave GDP-mannose, suggesting that SidI is a mannosyltransferase. However, future investigation is required to reveal the target of SidI’s activity *in vivo* and the molecular mechanism by which SidI inhibits translation.

SidI induces the host stress response through formation of a complex between heat shock factor 1 (HSF1) and eEF1A(12). The formation of this complex in conjunction with a non-coding RNA promotes the binding of HSF1 to the heat shock element (HSE), inducing host cell *Hsp70* expression (12, 36). Notably, despite the upregulation of transcription, Hsp70 protein levels did not differ between infected and uninfected cells (12), likely due to robust translation inhibition by *L. pneumophila* effectors. A more recent study revealed that the eEF1A1 isoform facilitates expression of heat shock genes independently of its role in protein translation (37). Since SidI, but not Lgt1, induced expression of *Hsp70* in addition to eEF1A-mediated activation of the heat shock factor 1 (HSF1) transcription factor, it could be hypothesized that SidI modulates eEF1A to specifically amplify heat shock genes, which could enhance survival of *L. pneumophila* infected cells. Moreover, the effector LegK4 phosphorylates Hsp70, which results in loss of protein translation. It is tempting to envision a scenario whereby SidI and LegK4 function in concert to modulate host Hsp70 activity. The importance for modulation of heat shock proteins has also been demonstrated by previous observations that Hsp90 is essential for *L. pneumophila* replication in *Acanthamoeba castellanii* (38). Further investigation is required to define the influence of SusF on SidI-mediated modulation of the heat shock response.

Regulation of effector function by metaeffectors is an essential component of the *L. pneumophila* virulence strategy. The first identified *L. pneumophila* metaeffector, LubX, functions as an E3 ubiquitin ligase that is translocated into the host cytosol at late stages of infection and catalyzes ubiquitination of the effector protein SidH, targeting it for proteasomal degradation (5). *L. pneumophila* Δ*lubX* mutants replicate similarly to wild-type bacteria, suggesting that LubX is not individually required for intracellular replication in host cells examined (5). Like SusF, the metaeffector SidJ directly contributes to intracellular replication by suppressing the toxicity of the SidE family of effectors (SdeA, SidE, SdeB and SdeC) (7, 39). However, unlike SusF, overexpression of SidJ is toxic to eukaryotic cells (39). The SidE family of effectors also contributes directly to intracellular replication through a novel mechanism of phosphoribosyl-ubiquitin conjugation to host substrates (40–42). Unlike the SidE effectors, loss of *sidI* or the *sidI*-*lpg2505* operon has no effect on *L. pneumophila* intracellular growth (6), suggesting that SidI functions redundantly within macrophages. However, in the absence of SusF, SidI is deleterious to *L. pneumophila* intracellular replication (6), a phenotype not observed for any other effector.

Several modes of metaeffector function have been described. A large-scale screen by Urbanus and colleagues led to the discovery of 17 novel *L. pneumophila* metaeffectors that function to suppress the toxicity of their cognate effectors (4). They uncovered several mechanisms by which metaeffectors regulate their cognate effectors through abrogation of effector enzymatic activity *in vitro* (LegL1, SidP, LupA). Similarly, we found that SusF dampens SidI activity *in vitro*; however, residual SidI activity is retained, suggesting that SusF functions to fine-tune SidI activity. Although we have not identified the *bona fide* substrate of SidI, our observation that SidI interacts with both SusF and eEF1A simultaneously at distinct regions suggests that SidI may modify eEF1A or use interaction with eEF1A to gain access to a host substrate. Future investigation will reveal the role of SusF in SidI function and how this contributes to *L. pneumophila* intracellular replication.

Through structural homology prediction and biochemical analysis, we revealed that SidI likely possesses GDP-dependent glycosyltransferase activity. Although we have not directly demonstrated the transfer of mannose to a substrate protein, it is unlikely that SidI functions only as a nucleotide-sugar hydrolase in the context of infection. Moreover, detection of free nucleotide liberation from nucleotide-sugar donors is a pre-requisite for transfer of glycans to substrate molecules and is an established method to identify glycosyltransferase activity without knowing the acceptor substrate (43). Our use of a luciferase-based assay eliminates the requirement for radioactive nucleotide-sugars.

Several *L. pneumophila* effectors function as glycosyltransferases, including the translation inhibitors Lgt1-3; however, none have been shown to utilize GDP-conjugated sugars. In fact, bacterial GDP-dependent glycosyltransferases seem to be involved primarily in biogenesis of cell surface structures (44–46). SidI has predicted structural homology to mycobacterial phosphatidylinositol mannosides (PIMs), which are GT-B glycosyltransferases (45, 47). To our knowledge, no other translocated bacterial effector has been demonstrated to have GDP-dependent glycosyltransferase activity. The primary site of protein glycosylation in eukaryotic cells is the Golgi apparatus; however, nucleotide sugars, including GDP-mannose, are synthesized in the cytoplasm of eukaryotic cells prior to transport into the Golgi (48, 49). Thus, SidI likely hijacks cytoplasmic GDP-mannose prior to its translocation into the Golgi. Based on the relatively low abundance of individual effectors in *L. pneumophila* infected cells it is unlikely that SidI function influences glycoprotein production by host cells. Thus, SidI is a GDP-dependent glycosyltransferase and this activity is likely critical for SidI-mediated translation inhibition and cytotoxicity. Further biochemical and structure-function analysis will reveal the detailed molecular mechanism by which SidI functions, its *bona fide* substrate in host cells and how SusF regulates its activity to promote *L. pneumophila* intracellular replication.

In this study, we have revealed a direct high-affinity interaction between SidI and its metaeffector SusF, defined a novel enzymatic activity for SidI and uncovered that SusF can modulate SidI function *in vitro*. Our work has provided a foundation for future biochemical and cell biological studies to reveal how SidI functions to modulate host cell processes. The severe virulence defect resulting from expression of *sidI* in the absence of *susF* underlies the importance of defining the molecular mechanism of SidI-SusF function. Moreover, the lack of precedent for a bacterial GDP-dependent glycosyltransferase effector protein suggests that SidI possesses novel enzymatic activity, which must be regulated by SusF to promote *L. pneumophila* intracellular replication. Our future work will focus on uncovering this mechanism in order to gain critical insight into translation inhibition and metaeffector function.

## Materials and Methods

### Bacterial strains, cell culture, growth conditions and reagents

*Escherichia coli* stains used for cloning (Top10; Invitrogen) and protein expression [BL21 (DE3); a gift from Dr. Craig Roy, Yale University] were maintained in Luria-Bertani (LB) medium supplemented with antibiotics as appropriate for plasmid selection [50 µg mL^-1^ kanamycin (GoldBio), 100 µg mL^-1^ ampicillin (GoldBio) and 25 µg mL^-1^ chloramphenicol (GoldBio)]. *Legionella pneumophila* Philadelphia-1 Lp01 (50) and Δ*dotA* (51) strains were cultured on supplemented charcoal–N (2-acetamido)-2-aminoethanesulfonic acid (ACES)-buffered yeast extract (CYE) and grown at 37°C as described previously (52, 53). CYE was supplemented with 10 µg mL^-1^ chloramphenicol for plasmid maintenance as required. Protein expression in *E. coli* and *L. pneumophila* was induced with 1 mM isopropyl-*β*-D-1-thiogalactopyranoside (IPTG) (GoldBio).

All mammalian cells were grown at 37°C/5% CO2 for up to 30 passages. HEK 293 cells were cultured in Dulbecco’s Modified Eagle Medium (DMEM; Gibco) supplemented with 10% heat-inactivated fetal bovine serum (HIFBS; Gibco). CHO FcγrII cells (54) (a gift from Dr. Craig Roy) were cultured in MEM*α* (Gibco) supplemented with 10% HIFBS.

Unless otherwise specified, all chemicals were obtained from MilliporeSigma (St Louis, MO). Oligonucleotide primers used in this study are listed in **Table 1**.

**Table 1.**
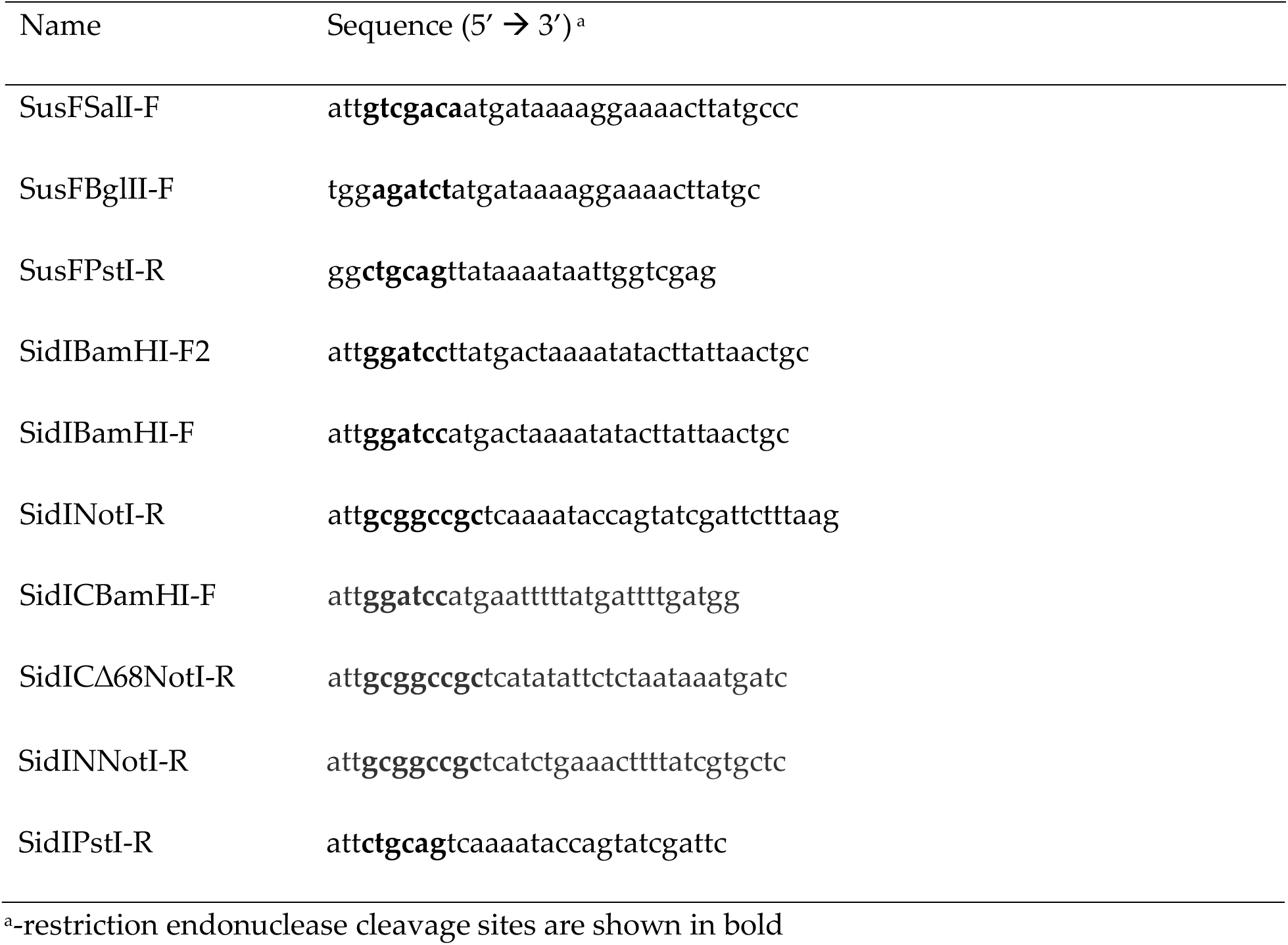
Oligonucleotide primers used in this study Name Sequence (5’ à 3’) ^a^

### Molecular cloning, plasmid construction and generation of *Legionella* strains

*Legionella pneumophila* Lp01 gDNA was isolated using the Illustra genomicPREP DNA isolation spin kit (GE Healthcare) and was used as a template for cloning *sidI* and *susF* into the indicated plasmid vectors. For recombinant protein production, *sidI* was amplified using primer pairs SidIBamHI-F/SidINotI-F and cloned as a BamHI/NotI fragment into pGEX-6P-1 (GE Healthcare) and pT7HMT (55). *susF* was amplified using primer pairs SusFBglII-F/SusFNotI-R and cloned as a BglII/NotI fragment into BamHI/NotI-digested pGEX-6P-1 and pcDNA4T/O-3xFLAG (56). For cloning into pT7HMT, *susF* was amplified using SusFSalI-F/SusFNotI-R primer pairs and cloned as a SalI/NotI fragment. For generation of SidI truncations, the regions of interest were amplified with primer pairs SidIBamHI-F/SidINNotI-R (SidIN), SidICBamHI-F/SidINotI-R (SidIC), or SidICBamHI-F/SidICΔ68NotI-R (SidICΔ68) and cloned as BamHI/NotI fragments into pT7HMT or pGEX-6P-1. For generation of p*cyaA*::*sidIRP*, *sidIR453P* was amplified from pSR47S::*sidIR453P* (6) with SidIBamHI-F2/SidIPstI-R and cloned as a BamHI/PstI fragment into p*cyaA* (21). DNA sequences were confirmed by Sanger sequencing (GENEWIZ, South Plainfield, NJ).

*L. pneumophila* Lp01 wild-type and Δ*dotA* producing CyaA-SidI constructs were generated by electroporation of p*cyaA*::*sidIRP* into competent strains using a BioRad Gene Pulser at 2.4 kV, 200 Ω, and 0.25 µF and plated on CYE supplemented with 10 µg mL^-1^ chloramphenicol. CyaA-SidI production was confirmed by Western blot as described below. *L. pneumophila* strains harboring p*cyaA*::*ralF* (21) were a gift from Dr. Craig Roy (Yale University).

### Transfection and selection of stable tissue culture cells

HEK 293 cells were transfected with pcDNA4T/O-3FLAG::*susF* using FuGENE HD (Roche) transfection reagent according to manufacturers’ instructions. At 48 h post-transfection, cells 500 µg mL^-1^ zeocin (Invitrogen) was added to culture medium and this selection was maintained for 10 days. Subsequently, cells were maintained in 200 µg mL^-1^ zeocin and production of 3FLAG-SusF was confirmed by Western blotting, as described.

### Recombinant protein expression and purification

Overnight *E. coli* BL21 (DE3) cultures were sub-cultured at 1:100 for 3 h in LB supplemented with the appropriate antibiotics followed by induction of protein expression with 1 mM IPTG, and induced at 16°C overnight. Bacterial cultures were centrifuged at 4,200 r.c.f. for 5 min at 4°C and washed with ice-cold PBS followed by incubation in bacterial lysis buffer [50 mM Tris pH 8, 100 mM NaCl, 1 mM ethylenediaminetetraacetic acid (EDTA), 200 µg mL^-1^ lysozyme, 2 mM dithiothreitol (DTT), 10 µg mL^-1^ DNase, and complete protease inhibitor] for 30 min on ice. Bacteria were sonicated on ice followed by centrifugation at 17,000 r.c.f. for 30 min at 4°C. Clarified bacterial lysates were incubated with either Ni-NTA beads plus 15 mM imidazole (His-tag) or Glutathione Agarose beads (GST-tag) for 1-2 h at 4°C with rotation. For His-tagged proteins, Ni-NTA agarose beads were transferred to 10 mL polyprep chromatography columns (BioRad) and washed with 20 mL of ice-cold wash buffer (50 mM Tris pH 8, 100 mM NaCl, 1 mM EDTA, 40 mM imidazole) followed by elution in 7 mL of elution buffer (50 mM Tris pH 8, 100 mM NaCl, 1 mM EDTA, 200 mM imidazole). For GST-tagged proteins, glutathione agarose beads were transferred to a 10 mL polyprep chromatography column and washed with 20 mL wash buffer I (PBS, 0.05% Triton X-100), followed by wash buffer II (PBS, 0.05% Triton X-100, 0.5 M NaCl) and eluted in GST elution buffer (50 mM Tris pH 9.5, 10 mM glutathione). Protein eluates were visualized by SDS-PAGE and Coomassie brilliant blue staining. Elution fractions containing protein were combined, dialyzed into PBS and quantified by Bradford Assay (Thermo Scientific).

In some cases, the poly-histidine tag was removed from recombinant proteins by site-specific proteolysis using Tobacco Etch Virus protease. Following digestion as previously described (55), the sample was reapplied to a Ni-NTA beads and the unbound fraction containing the target protein was collected. Samples were further purified by gel filtration chromatography using either a Superdex 75 (26/60) or Superdex 200 (26/60) column attached to an AKTA-format FPLC (GE Healthcare) and PBS as a running buffer. Elution fractions containing protein were analyzed by SDS-PAGE and Coomassie brilliant blue staining, as described above.

### Affinity chromatography for protein-protein interaction

Overnight *E. coli* cultures grown in LB plus appropriate antibiotic were sub-cultured 1:100 into 20 mL LB and grown at 37°C for 3 h followed by induction with 1 mM IPTG for 4 h. Cultures were centrifuged for 10 min at 1500 r.c.f., washed with ice-cold PBS and pelleted in 1 mL aliquots prior to storage at −20°C for <72 hours until use. Pellets were re-suspended in 1 mL of cold lysis buffer (50 mM Tris pH 8, 100 mM NaCl, 1 mM EDTA) supplemented with 200 µg mL^-1^ lysozyme, and complete protease inhibitor and incubated on ice for 30 min. Two millimolar DTT was added to lysates before sonicating on ice. Lysates were then clarified by centrifugation at 12,600 r.c.f. for 15 min at 4°C. Supernatants were collected in fresh, pre-chilled microcentrifuge tubes. Supernatants were added to either pre-equilibrated Ni-NTA magnetic beads or magnetic glutathione agarose beads following manufacturer’s protocol (Pierce) for binding His- or GST-tagged fusion proteins, respectively. After the initial batch binding, beads were washed 2x with wash buffer [Ni-NTA: 50 mM Na3PO4, 300 mM NaCl, 15 mM imidazole, 0.05% Tween-20, pH 8; Glutathione: 125 mM Tris-Cl, 150 mM NaCl, 1mM DTT, 1 mM EDTA, pH 7.4] prior to addition of recombinant protein or subsequent cell lysate and batch binding for 1 h at 4°C with rotation. Beads were washed 2x with wash buffer before adding the final cell lysate or recombinant protein and incubating for 1 h at 4°C with rotation. Following binding, beads were washed 2x with wash buffer and transferred to fresh, pre-chilled 1.5 mL microcentrifuge tubes and resuspended in 25 µL of 3x Laemmli sample buffer and boiled for 10 min. Samples were analyzed by SDS-PAGE and Coomassie brilliant blue or Western Blot, as indicated.

For experiments to examine eEF1A binding, lysates were derived from either HEK293T or HEK293 3FLAG-SusF cells grown to ≥70% confluence on tissue culture (TC) treated dishes. Cells were washed 1x with ice-cold PBS and lysed in 1 mL of mammalian lysis buffer [1% NP-40 (v/v), 150 mM NaCl, 20 mM Tris-Cl pH 7.5, 10 mM Na4P2O7, 50 mM NaF, and complete protease inhibitor]. Lysates were collected in pre-chilled, 1.5 mL centrifuge tubes and centrifuged at 11,000 r.c.f. for 20 min at 4°C. Supernatants were collected in pre-chilled, 1.5 mL centrifuge tubes and stored on ice until use.

### CyaA effector translocation assay

Translocation of CyaA fusion proteins was performed as described (21). Briefly, CHO FcγrII cells (a gift from Dr. Craig Roy) were seeded into 24-well plates at 1×10^5^ cells per well 24 h prior to infection. Cells were infected at a multiplicity of infection (MOI) of 30 with *L. pneumophila* Lp01 wild-type or Δ*dotA* that had been cultured on CYE supplemented with 10 µg mL-1 chloramphenicol and 1 mM IPTG followed by opsonization by incubation for 30 min with an *α*-*L. pneumophila* antibody (Invitrogen, PA17227; 1:1000) at RT. Cell culture media was supplemented with 1 mM IPTG to ensure expression of CyaA-SidI fusion protein. Infected cells were incubated for 1-2 hours at 37°C/5% CO2 followed by aspiration of culture media and washing 3 times with ice-cold PBS. Cells were lysed for 30 minutes in 200 µL of ice-cold lysis buffer (50 mM HCl, 0.1% Triton-X100) with rocking at 4°C. Samples were either stored at −80°C until use or immediately mixed with 12 µL 0.5M NaOH in 95% ethanol and centrifuged for 5 min at 11,000 r.c.f. Supernatants were dried in a Speed Vac and stored at −80°C until use. cAMP was quantified using a cAMP Direct Biotrak EIA (non-acetylation) kit (GE Healthcare). Briefly, dried samples were resuspended in 250 µL Assay Buffer and ELISA was performed following manufacturer’s protocol. Absorbance at 655 nm was read in a Victor 2 microplate reader (PerkinElmer).

### *In vitro* protein translation assay

Translation assays were performed using the Promega Flexi^®^ Rabbit Reticulocyte Lysate System (RLL; L4540) following manufacturer’s protocol. Briefly, a master mix of RLL was generated and aliquoted into individual tubes. Four nanograms of purified His6-SidI (see above) were added as indicated. Purified recombinant (untagged) SusF was added at the indicated concentrations. Proteins were equilibrated to room temperature before use. All reactions were brought to 50 µL with ultra-pure water. Reactions were mixed by pipetting and briefly centrifuged before incubation at 30°C for 90 min. Translation of *Firefly* luciferase mRNA (Promega) was quantified using a Victor 2 microplate reader (PerkinElmer).

### Glycosyltransferase activity assay

Glycosyltransferase activity was evaluated using GDP- or UDP-Glo^TM^ Glycosyltransferase Assay kits (Promega) with GDP-mannose (VA1095) or UDP-glucose (V6991), respectively, following manufacturers’ recommendations. Briefly, 5 µg of purified His_6_-SidI and/or molar equivalent (or excess as indicated) SusF were added to 100 µL of 50mM Tris pH 7.4 with 10 µM of GDP-mannose or UDP-glucose. Ten µM GDP or UDP were used as controls as indicated. Reactions were carried out for an hour at 37°C and quantification of free nucleotide (GDP or UDP) was achieved by addition of GDP- or UDP-Glo^TM^ nucleotide detection reagent following manufacturers’ instructions and analyzed via luminescence using a Victor 2 microplate reader (PerkinElmer).

### SDS-PAGE and Western Blot

Boiled protein samples were loaded onto either 4-20% gradient SDS-PAGE gels (BioRad), 12% or 15% SDS-PAGE gels. Following electrophoresis, proteins were visualized with Coomassie brilliant blue or transferred to PVDF membranes using a BioRad wet transfer cell. Membranes were incubated with blocking buffer [5% nonfat milk powder dissolved in tris-buffered saline- 0.1% Tween 20 (TBST)]. Primary antibodies [*α*-eEF1A (#2551S Cell Signaling Technology), *α*-Flag-M2 (Sigma), or *α*-CyaA[3D1] (#EG800 Kerafast)] were used 1:1000 in blocking buffer and detected with HRP-conjugated secondary antibodies (1:5000; ThermoFisher). Membranes were washed, incubated with ECL substrate (GE Amersham) and imaged by chemiluminescence using an Azure Biosystems c300 Darkroom Replacer.

### Surface Plasmon Resonance

Direct binding of His_6_-SidI to SusF was assessed by SPR using a Biacore T-200 instrument (GE Healthcare) at 25 °C, according to the general methods previously described (57). Briefly, all experiments were carried out in a running buffer of HBS-T (20 mM HEPES (pH 7.4), 140 mM NaCl, and 0.005% (v/v) Tween-20 and a flow-rate of 30 µl/min (57). SusF (50 µg mL^-1^ in 10 mM acetate, pH 4.5) was immobilized to a final density of 1063 RU on one flow cell of a CMD-200M surface (XanTec Bioanalytics, GmbH; Dusseldorf, Germany) using standard NHS/EDC coupling. A reference surface was prepared in a similar manner by ethanolamine quenching of NHS/EDC-activated flow cell. Experimental sensorgrams of His_6_-SidI binding to immobilized SusF were obtained in reference-corrected, single-cycle mode using sequential concentrations of 0.8, 4, 20, 100, and 500 nM His_6_-SidI. Each association phase consisted of 2 min sample injection, followed by a 1.5 min dissociation phase, except for the final injection which incorporated a 60 min dissociation phase for more accurate determination of the dissociation rate. Kinetic analysis was performed using Biacore T-200 Evaluation Software v3.1 (GE Healthcare). Sensorgrams were analyzed using both Langmuir and Two-State Reaction binding models and a local value of R_max_.

### Analytical Gel Filtration Chromatography

Samples of recombinant proteins were characterized by analytical-scale gel-filtration chromatography as a mean of assessing their apparent molecular weight. All samples and standards (500 µl total volume) were separated on a Superdex 200 10/300 column (GE Healthcare) attached to an AKTA-format FPLC system using a flow-rate of 0.5 ml/min and PBS as a running buffer. Fractions of 1 ml were collected for subsequent analysis by SDS-PAGE and Coomassie brilliant blue, as described above.

### Molecular Modeling

Homology models were created using four different servers, using the full sequence of SidI as inputs and default parameters for each server. Using HHPred (23), a number of templates were identified for residues 376 to 868, and the top 25 templates were forwarded for modeling. Using Phyre2 (24), a smaller region, residues 585 to 763, was chosen for modeling. Raptor-X (20) and I-TASSER (25–27) both produce full-length models. Raptor-X predicted residues 350 to 870 to be a domain. I-TASSER presented five models of low confidence, one of which contained a glycosyltransferase domain with a GT-B fold, which was selected as the working model.

### Statistical Analysis

Statistical analysis was performed with GraphPad Prism software using Students’ t-test, as indicated, with a 95% confidence interval. For all experiments, data are expressed as mean ± standard deviation (s.d.) of samples in triplicates.

## Acknowledgements

We thank Drs. Philip Hardwidge and Mary Weber for critical reading of the manuscript and Dr. Craig Roy for providing strains and cell lines. This work was funded through an NIH NIGMS COBRE Research Project Award (P20GM113117; to SRS), a Kansas-INBRE Developmental Research Project Award (P20GM103418; to SRS), a Kansas State University College of Arts and Sciences Undergraduate Research Award (to TJB), a Kansas-INBRE Semester Scholar Award (P20GM103418; to AEP), and start-up funds from Kansas State University (to SRS).

## Supplemental Information

### Supplemental Figure Legends

**Figure S1.**
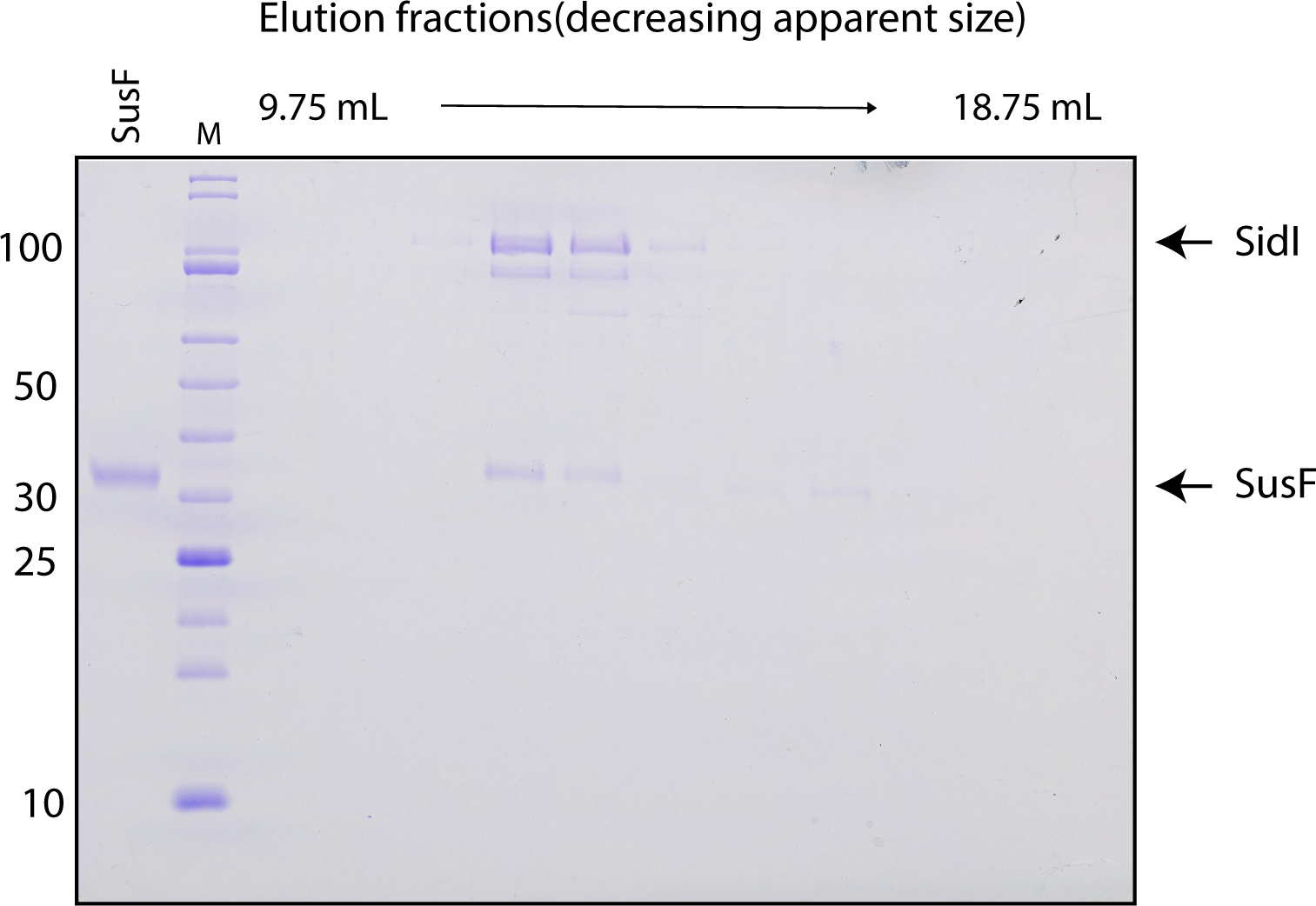
Co-elution of SusF and SidI by gel filtration chromatography. A sample of recombinant SidI bound to SusF was separated by analytical scale gel filtration chromatography as presented in Figure 1C. Ten µl of each column fraction were analyzed by SDS-PAGE followed by Coomassie staining to assess the protein content within each fraction. Fractions corresponding to the peak of approximately 150 kDa apparent molecular weight in the sample of SidI bound to SusT (Fig. 1C, red trace) contained both proteins, consistent with formation of a 1:1 complex between these two molecules.

**Figure S2.**
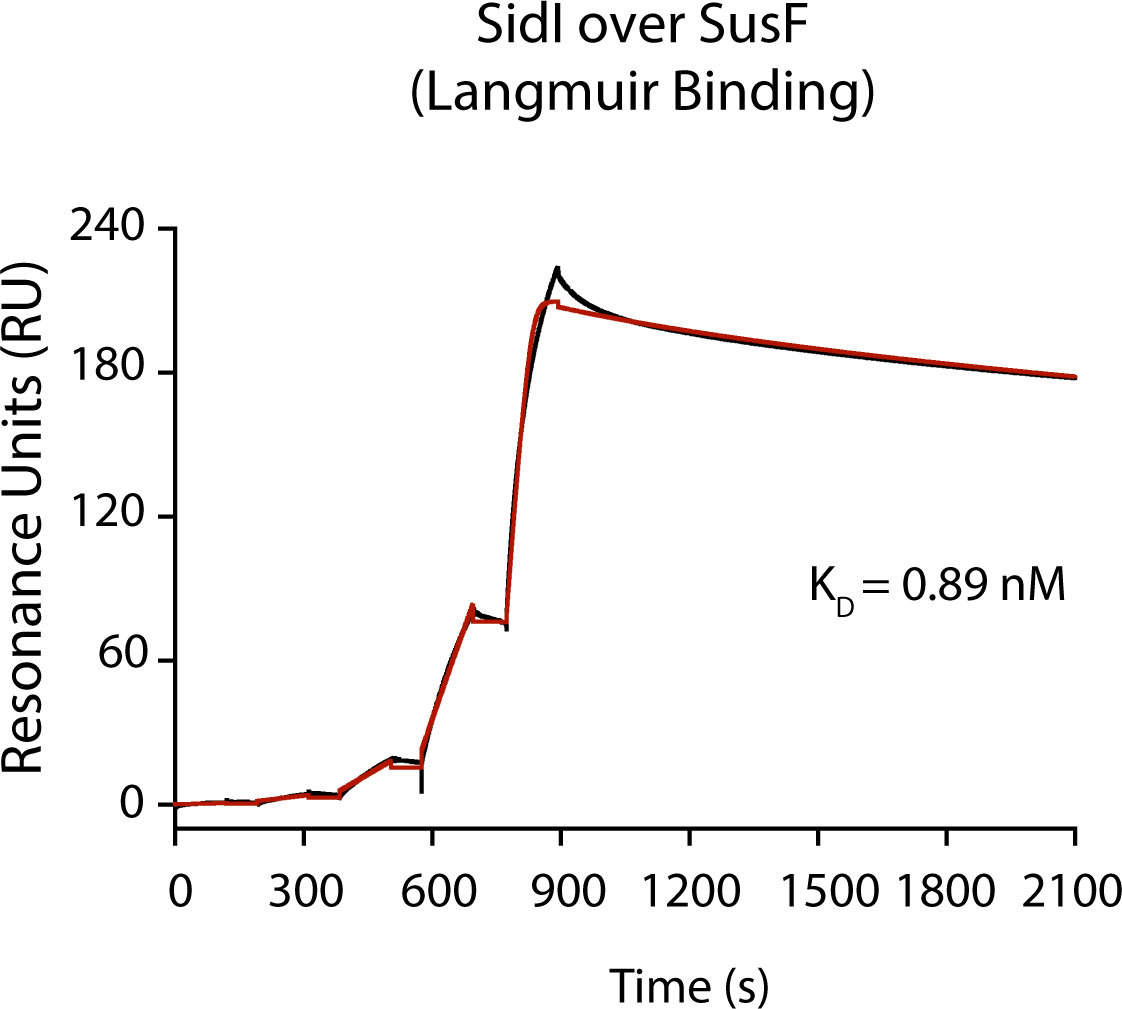
Binding of His6-SidI to immobilized SusF was assessed by SPR. The reference corrected sensorgram from a single-cycle experiment is shown in black, while the outcome of fitting to a Langmuir binding model is shown in red. Using this model, the interaction is described by an apparent KD of 0.89 nM where kon = 2.2×10^5^ M^-1^s^-1^ and koff = 1.9×10^-4^ s^-1^, respectively.

**Figure S3.**
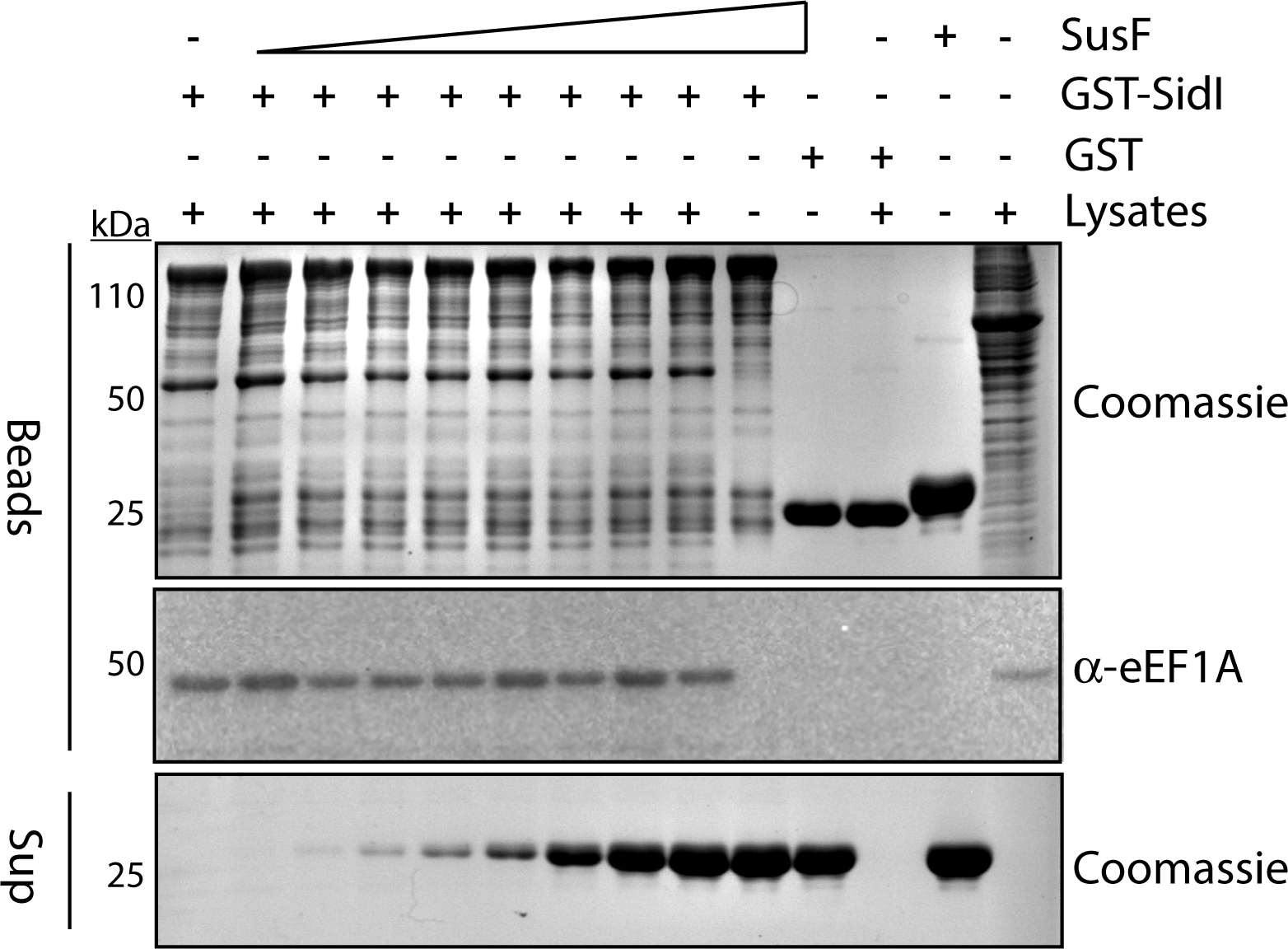
eEF1A interacts with the SidI-SusF complex. Lysates from *E. coli* expressing GST-SidI or GST alone were incubated with magnetic glutathione agarose beads and washed followed by addition of 10, 25, 50, 100, 200, 400, 800 or 1000 µg of purified recombinant SusF (shown as increasing amounts in supernatants from beads). Beads were subsequently incubated with lysates from HEK 293T cells. Proteins remaining on the beads were separated by SDS-PAGE and visualized by Coomassie stain or Western blot

**Figure S4.**
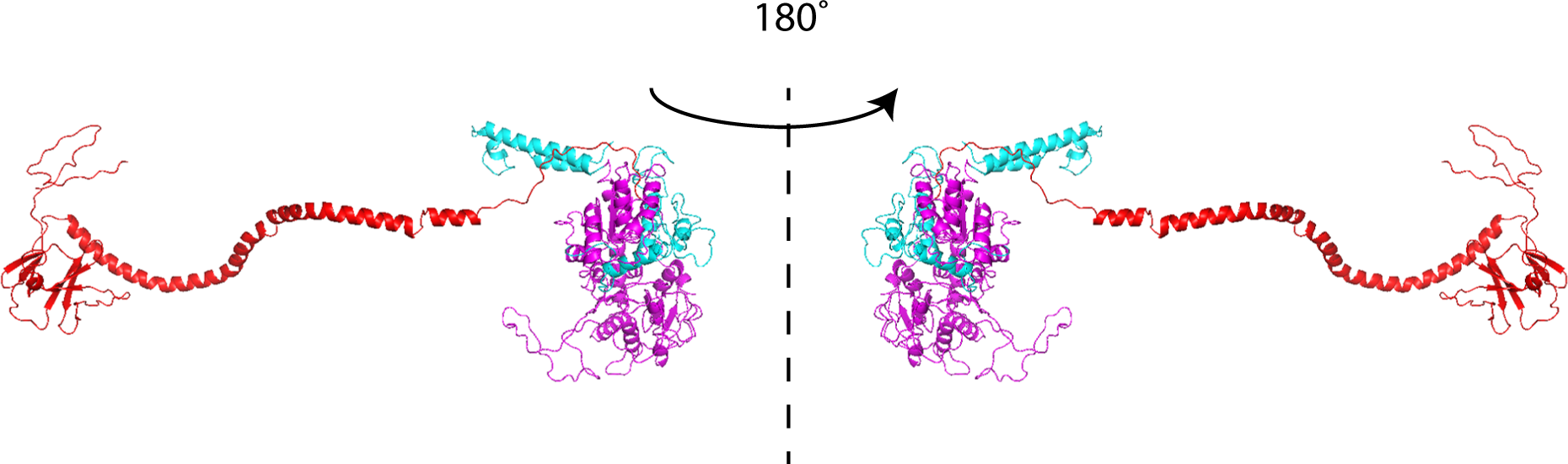
Molecular model of SidI. Ribbon cartoons of SidI modeled using the RaptorX webserver. Putative SidI domains are shown in red (residues 1-268), magenta (residues 269-874) and cyan (residues 874-942).

**Figure S5.**
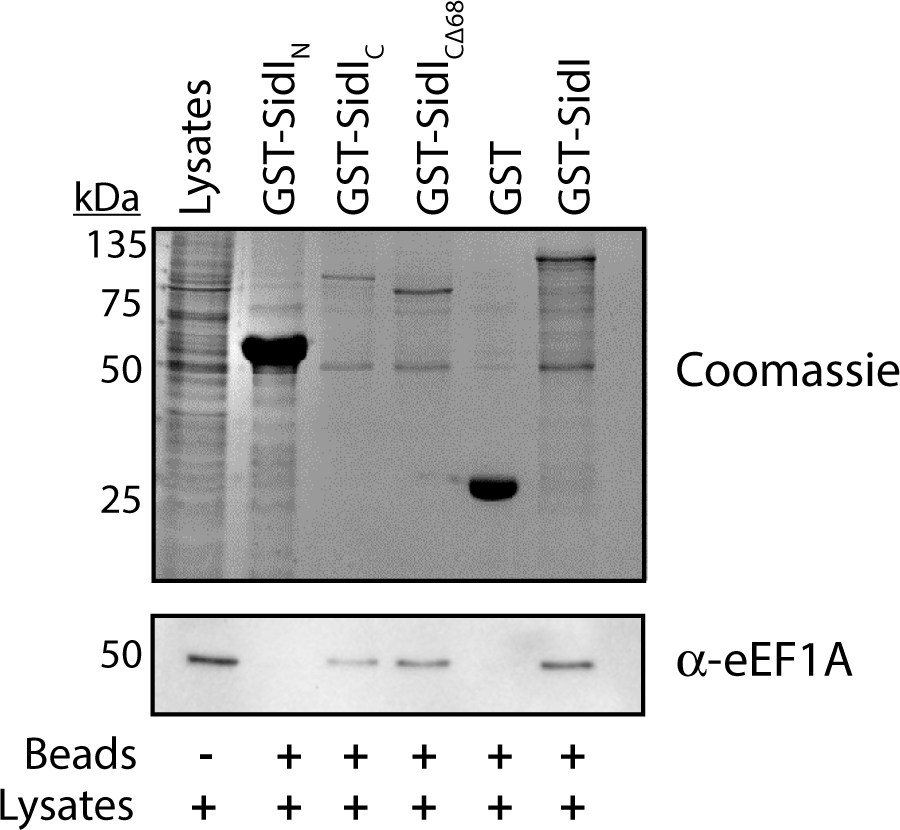
Interaction between eEF1A and SidI truncations in the absence of SusF. Lysates from *E. coli* expressing GST-SidI constructs were incubated with glutathione agarose beads followed by washing and incubation with lysates from HEK 293 cells. Proteins were separated by SDS-PAGE and visualized by Coomassie staining (GST-SidI) or Western blotting (eEF1A). Data are representative of two independent experiments.

**Figure S6.**
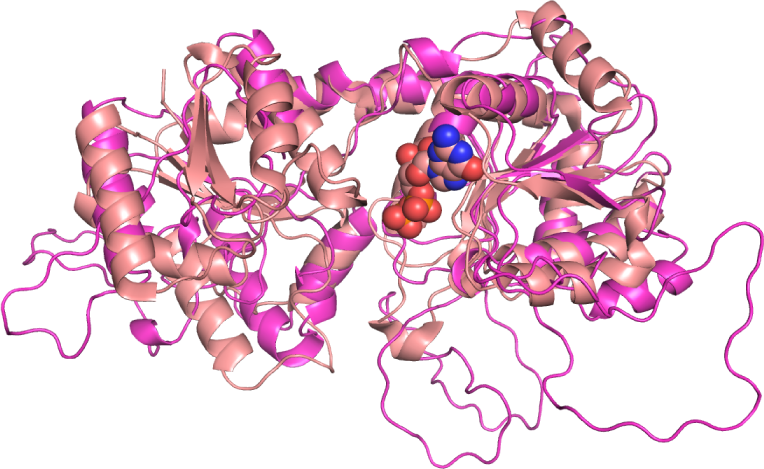
SidI is modeled as a glycosyltransferase with a GT-B fold. The model of the putative glycosyltransferase domain is presented in magenta ribbons superimposed to that of PimB (PDB: 3OKA) represented as salmon ribbons, which was identified by DALI as the most similar structure in the PDB. The structure of PimB was solved in the presence of GDP (salmon spheres).

### Supplemental Tables

**Table S1.**
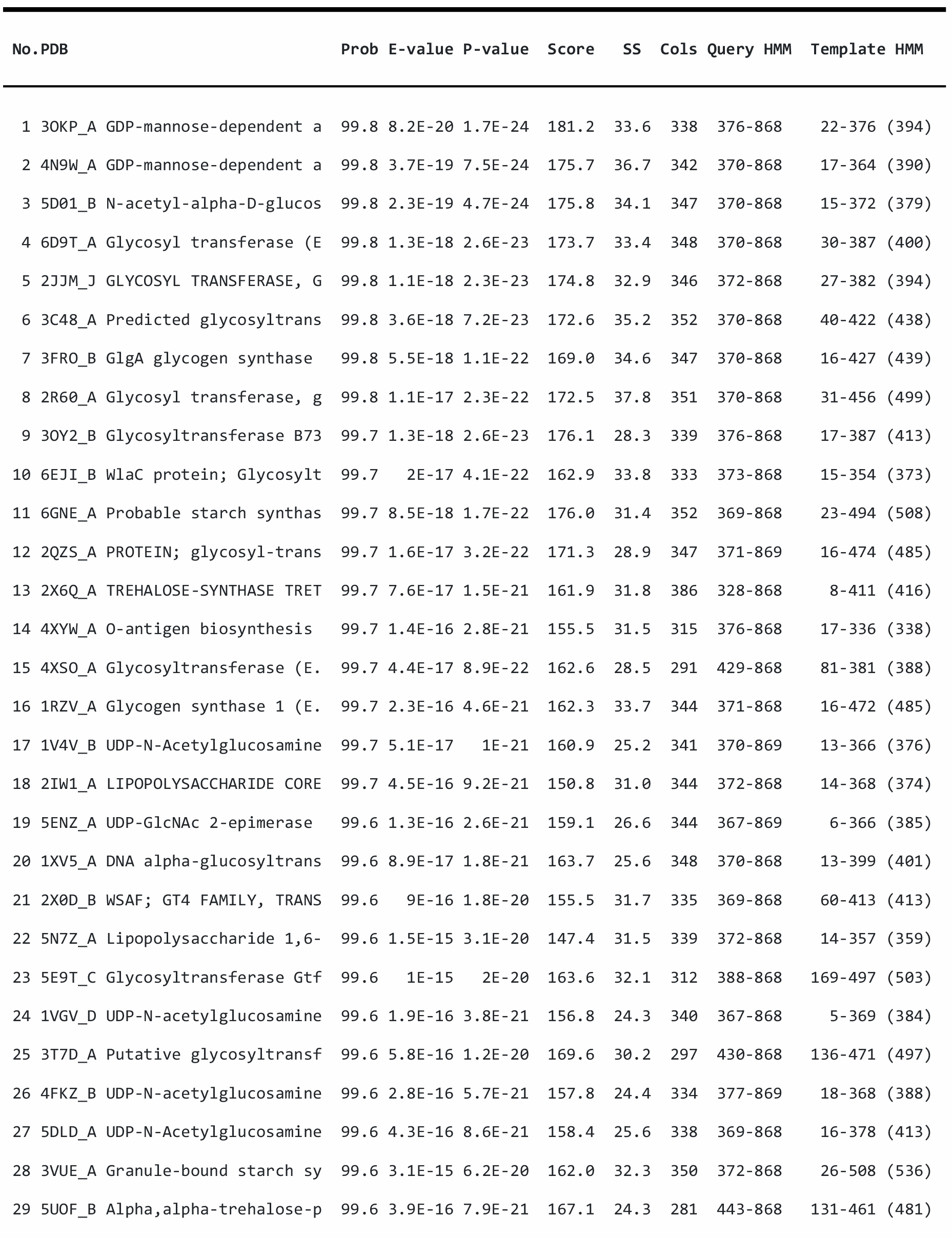
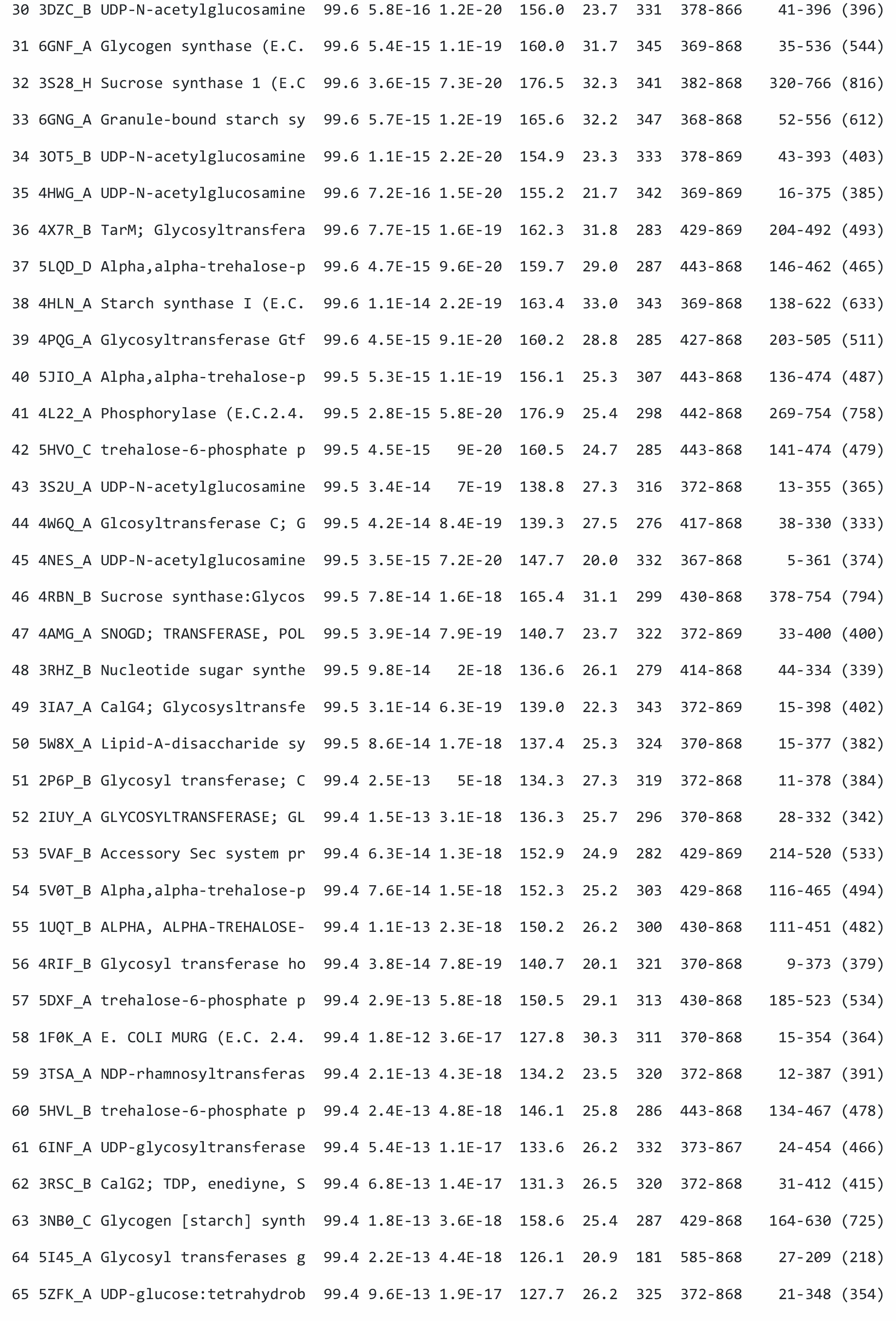
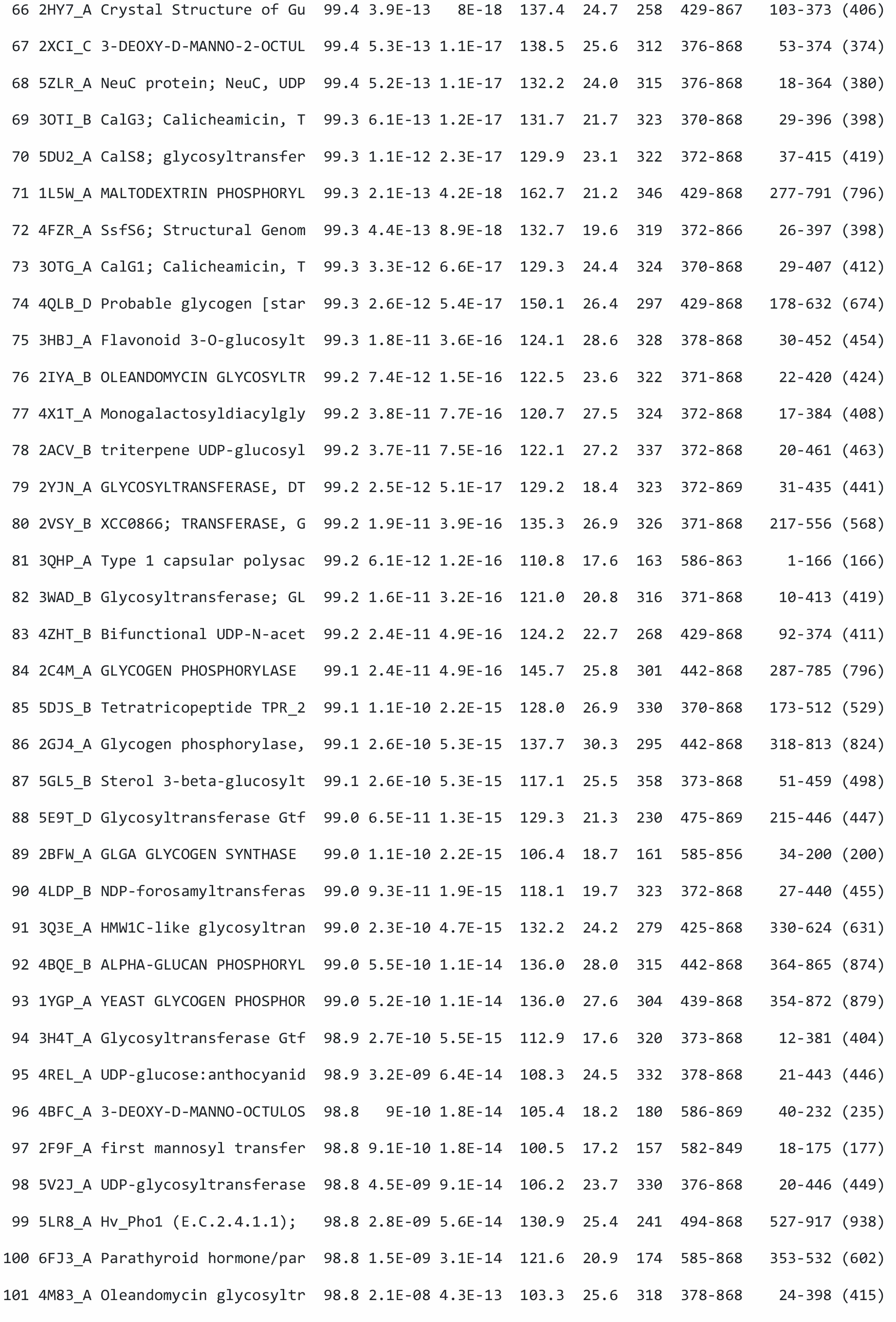
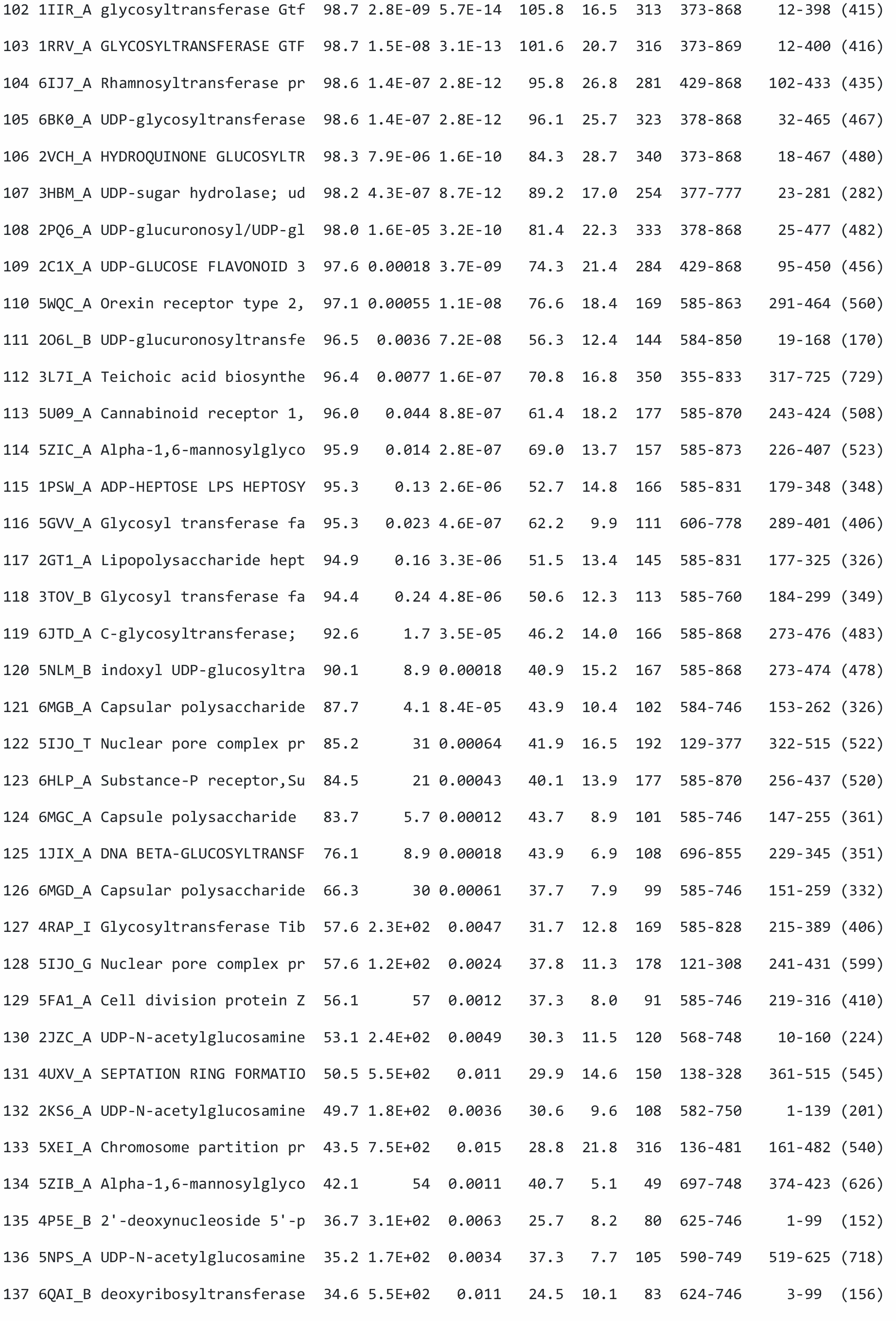
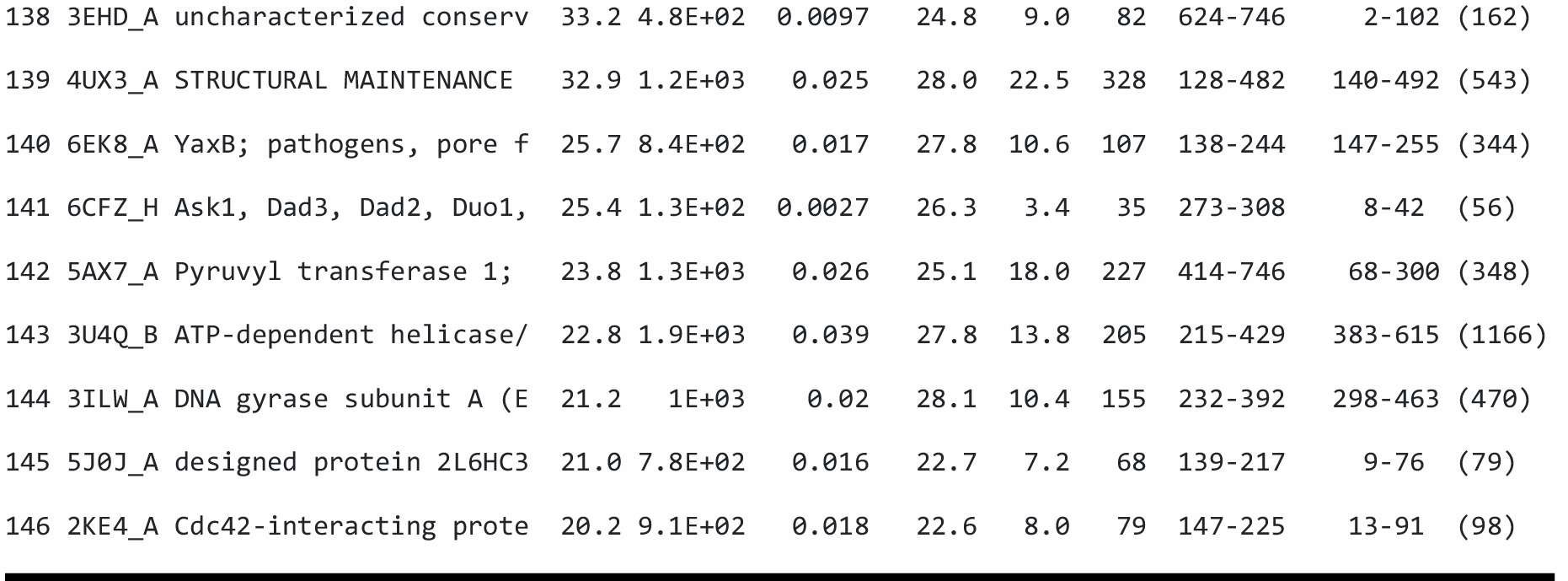
Proteins with homology to SidI (HHPred).

